# Genomic and experimental evidence that alternate transcription initiation of the Anaplastic Lymphoma Kinase (ALK) kinase domain does not predict single agent sensitivity to ALK inhibitors

**DOI:** 10.1101/696294

**Authors:** Haider Inam, Ivan Sokirniy, Yiyun Rao, Anushka Shah, Farnaz Naeemikia, Edward O’Brien, Cheng Dong, David McCandlish, Justin R Pritchard

## Abstract

Genomic data can facilitate personalized treatment decisions by enabling therapeutic hypotheses in individual patients. Conditional selection, which includes mutual exclusivity, is a signal that has been empirically useful for identifying mutations that may be sensitive to single agent targeted therapies. However, a low mutation frequency can underpower this signal for rare variants and prevent robust conclusions from genomic data. We develop a resampling based method for the direct pairwise comparison of conditional selection between sets of gene pairs. This effectively creates positive control guideposts of mutual exclusivity in known driver genes that normalizes differences in mutation abundance. We applied this method to a transcript variant of anaplastic lymphoma kinase (ALK) in melanoma, termed ALK^ATI^, which has been the subject of a recent controversy in the literature. We reproduced some of the original cell transformation experiments, performed rescue experiments, and analyzed drug response data to revisit the original ALK^ATI^ findings. We found that ALK^ATI^ is not as mutually exclusive with BRAF or NRAS as BRAF and NRAS genes are with each other. We performed *in vitro* transformation assays and rescue assays that suggested that alternative transcript initiation in ALK is not likely to be sufficient for cellular transformation or growth and it does not predict single agent therapeutic dependency. Our work strongly disfavors the role of ALK^ATI^ as a targetable oncogenic driver that might be sensitive to single agent ALK treatment. The progress of other experimental agents in late-stage melanoma and our experimental and computational re-analysis led us to conclude that further single agent testing of ALK inhibitors in patients with ALK^ATI^ should be limited to cases where no other treatment hypotheses can be identified.

## Introduction

In cancers, clonal selection influences tumor progression and responses to therapy (1), but in every patient, the process of clonal selection creates independent tumors with independent, parallel evolutionary trajectories. These parallel evolutionary paths can be conditioned upon the occurrence of a previous genetic event. Co-occurrence is when the first variant predicts the presence of a second (2,3). Whereas, mutual exclusivity occurs when the presence of the first predicts the absence of the second—and is the primary focus here. Mutual exclusivity is often viewed qualitatively or used to build large scale networks (2,4–6). Here we aim to create quantitative guideposts of conditional selection that will allow for direct comparisons of mutual exclusivity relative to known druggable oncogene pairs by controlling for cohort size. These guideposts can be used to triage the translational actionability of rare genomic findings.

A strong example of mutual exclusivity in cancer is seen in the evolutionarily ancient mitogen-activated protein kinase (MAPK) pathway that is present from yeast to metazoans (7). In higher eukaryotes, the MAPK pathway is often canonically activated by receptor tyrosine kinases (RTK) that signal through MAPKs to achieve cellular growth and development (8,9). Because of their critical role in growth and division, most known RTK-MAPK mutational events that drive cancer growth lead to mutual exclusivity across patients during parallel evolution. Moreover, many of the most impressive success stories for targeted cancer therapy in the past two decades have centered on this one pathway. Multiple examples of mutual exclusivity have been found between ALK-fusion, epidermal growth factor recepter (EGFR), Kristen rat sarcoma viral (KRAS), and Erb-b2 receptor tyrosine kinase 2 (ERBB2) genes in non-small cell lung cancer (NSCLC) (10–12). This has led to inhibitors of ALK, EGFR, and recently KRAS_G12C_ offering high rates of single agent therapeutic responses (13–17). In thyroid cancer, BRAF mutations have been found to be mutually exclusive with RET fusions, and both have shown high single agent response rates in clinical trials (18,19). Similarly, BRAF & NRAS are mutually exclusive in melanoma and single agent use of vemurafenib leads to clinical responses (20–26). Therefore, mutually exclusive activating mutations in the RTK-MAPK pathway have shown impressive clinical sensitivity to single agent targeted therapies (15,27).

There are at least two evolutionary explanations for mutual exclusivity in gain of function oncogenes in the same pathway: 1) Variants that are sufficient to activate the pathway may be functionally redundant. In this case there is no added benefit to a second activating mutation. 2)When activation of a single pathway member is sufficient, further activation of that pathway (via a second activating mutation) might be harmful to the cell i.e. the two events in the same pathway may be antagonistic because excess pathway activity is selected against (28,29).

While mutual exclusivity is often measured in cancer genomics studies, it can be challenging to interpret for rare genomic findings. For instance, KRAS and EGFR are well described as mutually exclusive in the literature, and their inhibitors have shown single agent response rates. If a newly discovered variant appears to be mutually exclusive with one or both of these genes, that would give confidence that the variant is potentially indicative of therapeutic response. However, when the variant is not mutually exclusive, the important question becomes, how do we demonstrate a negative result? Answering this question requires estimating the variability in effect size and significance that rarity would bring to positive control genes like KRAS and EGFR. In the absence of such a quantitative test, it is easy to view the data qualitatively (30). Here we seek a direct and quantitative test to rapidly interpret the relative mutual exclusivity of rare variants versus a positive control pair. Since we use positive controls that are genes in the MAPK pathway with previously verified single agent responses to targeted therapeutics (14–16,31), our method can triage rare genomic findings for potential single agent drug efficacy. If a newly discovered alteration is as mutually exclusive as a therapeutically established mutually exclusive gene pair, it gives greater confidence in the potential for single agent therapy. The decision tree in Supplementary Fig 1 illustrates the advances in mutual exclusivity analysis that are enabled by our method. To our knowledge, this is the first method that enables a quantitative and statistically robust analysis of a negative finding of conditional selection.

As a test case for a method that can quantitatively compare mutual exclusivity between gene pairs, we re-examined the observation that alternative transcription initiation in ALK, termed ALK^ATI^, is a therapeutically actionable oncogenic target (32). This novel ALK transcript has a transcription initiation site in intron 19 of ALK, just upstream of ALK’s kinase domain. Wiesner *et al* posited that this transcript exhibits a novel mechanism of oncogenic activation by overexpressing the kinase domain of ALK. Using *in vitro* transformation assays and inhibitor treatment, they hypothesize that single agent ALK-inhibitor therapy can treat the 2-11% (32,33) of melanoma patients expressing ALK^ATI^. However, Couts *et al* reported crizotinib (an ALK inhibitor) sensitivity in melanoma patient derived xenografts (PDXs) expressing EML4-ALK but not ALK^ATI^ *in vitro* and *in vivo* (34). Exome sequencing of the melanoma PDXs expressing ALK^ATI^ revealed that 4 out of 6 of these cell lines had transforming mutations in well-established melanoma oncogenes. Furthermore, out of the two ALK^ATI^ melanoma patients treated with ALK-inhibitors so far, one had a modest response that did not rise to the level of an objective response (32) and the other did not respond (34). This contradictory evidence, and the small sample size of the Couts *et al*. study suggest that reanalysis of the original observation of ALK^ATI^ is important to understanding if further investigation is warranted.

In this paper, we develop a simple and user-friendly resampling method to test the relative conditional selection across sets of gene pairs. This will allow us to quantitatively compare any gene of interest (GOI) such as ALK^ATI^ with positive control gene pairs like BRAF and NRAS in melanoma. Using this method, we show that ALK^ATI^ is not as mutually exclusive with BRAF or NRAS as BRAF and NRAS are with each other. Moreover, by large scale repetition of experiments performed in the Wiesner paper (32), we uncovered kinase activating mutations in ALK^ATI^ can be selected for during transformation assays. This suggests that it is not sufficient for growth factor independent transformation. We also find that ALK^ATI^ cannot compensate for oncogenic signaling in melanoma cells, and that ALK kinase domain overexpression does not predict ALK inhibitor sensitivity in the cancer cell line encyclopedia (CCLE) (35). Finally activated ALK actually inhibits the growth of BRAFV600E melanoma cell lines. Given the stunning advances in melanoma therapy, and the rich clinical trial landscape, we suggest that relapse/refractory melanoma patients with ALK^ATI^ should not be given ALK inhibitors in an investigational or off-label capacity unless no other options remain.

## Results

### Pairwise comparisons of conditional selection allow direct comparisons between frequent and rare events

Highly mutually exclusive gene pairs such as KRAS/EGFR and EGFR/EML4-ALK in lung cancer, and NRAS/BRAF in melanoma predict the success of ALK, EGFR and BRAF inhibition in the clinic (15,27). These mutually exclusive gene pairs are positive controls that can be compared with any gene/alteration of interest (GOI, in Fig 1A). A direct pairwise comparison would allow us to ask whether a “BRAF/NRAS level” of mutual exclusivity exists between any new GOI (such as ALK^ATI^) and BRAF or NRAS (BRAF and NRAS are an example of a highly mutually exclusive gene pair in melanoma, 20–24). However, the differences in sample size due to mutation frequency complicates our ability to directly compare the odds ratios between distinct gene pairs. If the GOI is much less abundant than the positive control, how often would the positive control have a similar odds ratio if it was only as abundant as the test gene of interest? Resampling will allow us to more directly compare the mutual exclusivities between pairs of genes with unequal abundances in cancer genomes (see method and supplement for full details).

**Figure 1:**
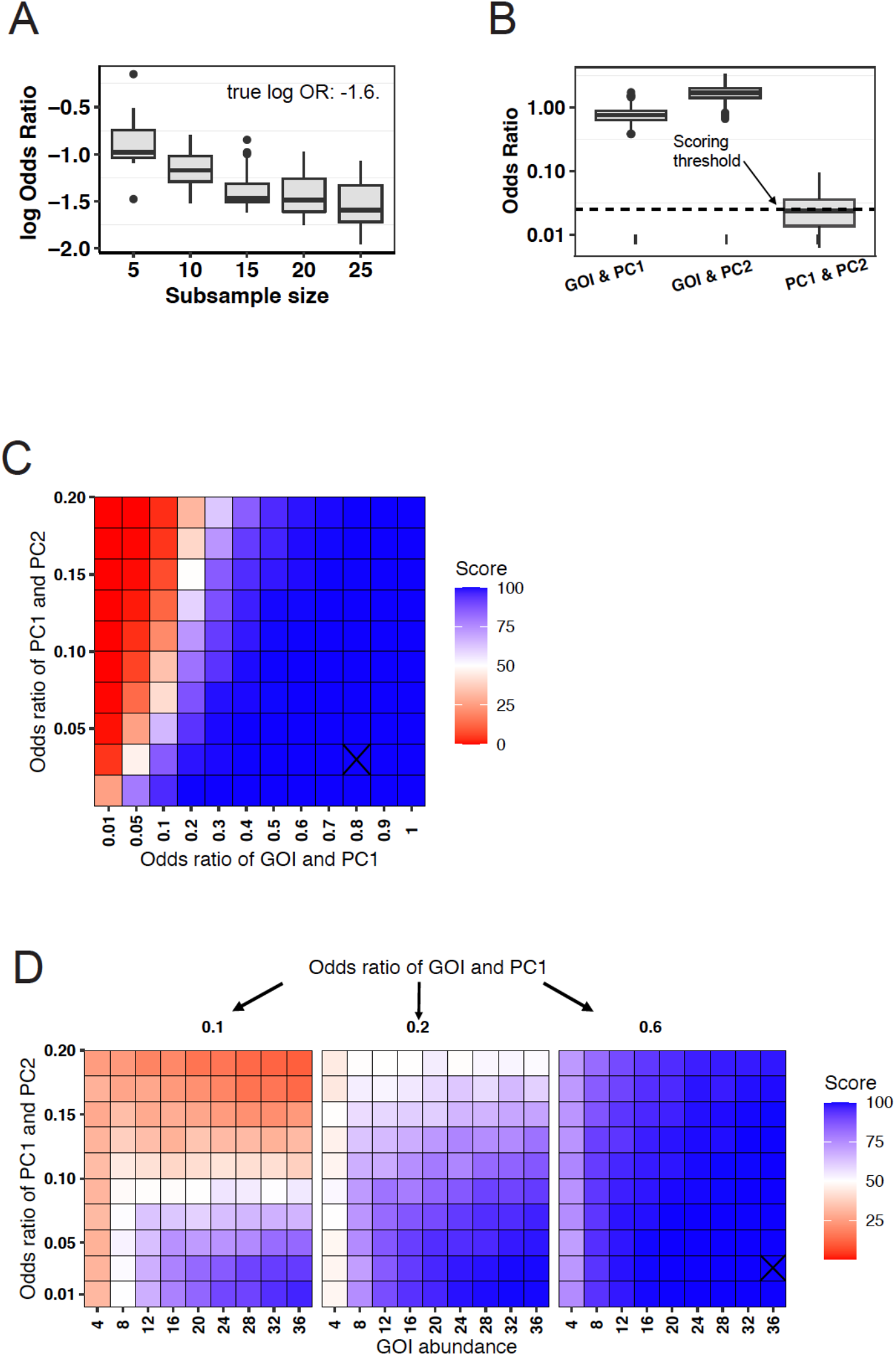
Pairwise comparisons of conditional selection between a GOI gene pair and a positive control gene pair in simulated cohorts. **(A)** Distribution of odds ratio of two mutually exclusive genes when one of the genes is rare (left-side) and abundant (right-side). **(B)** A single simulated patient cohort is scored by our pairwise comparisons approach. In this cohort, the odds ratio of PC1 with PC2 was 0.05 and the odds ratio of GOI with both PC1 and PC2 genes was >1. The frequency of GOI was set to 5%. The scoring threshold is the median of the PC1 and PC2 distribution. The score is the percentage of the GOI & PC1 or GOI & PC2 distribution that is greater than this threshold. Odds ratios displayed on the plot are for the comparison between the gene of interest and positive control (gene of interest gene pair). **(C-D)** Heatmaps showing the dependence of a high score (blue) on the difference between the odds ratios of the two gene pairs and on the abundance of the GOI. Each tile represents a single simulated cohort. A cohort size of 500 simulated patients was used. A GOI abundance of 25 patients was used in **C**.

To directly illustrate this idea, we analyzed how sample size affects the odds ratios observed for a simulated mutually exclusive positive control gene pair. We created a mock clinical cohort with two abundant but mutually exclusive genes (odds ratio of 0.025), and then we performed resampling experiments at different subsample sizes. As expected, decreasing the abundance of one of the genes increases the range in the observed odds ratios (OR) at smaller sample sizes: OR of 0-0.7 for a subsample size of 5 and OR of 0-0.11 for a subsample size of 20 (Fig 1A). Therefore, the range of odds ratios that is observable in highly mutually exclusive genes becomes noisier as sample size decreases. To more directly compare gene pairs with different abundances, we propose adjusting the frequency of the more prevalent genes to a frequency that is matched by the gene of interest.

We term this idea, “pairwise comparisons of conditional selection”. We aim to directly compare two gene pairs while controlling for the differences in sample size. Given two highly prevalent, mutually exclusive positive control genes and a rare GOI with low frequency, our method adjusts the frequency of one of the positive control genes to match frequency of the rare GOI. This effectively normalizes the prevalence of rare vs common genes. Comparing the odds ratios produced by the two positive control genes to the odds ratios produced by the GOI with one of the positive control genes lets us test the null hypothesis that no difference exists between the odds ratios of the two gene pairs (Fig 1B). Details of the pairwise comparison strategy are included in the methods section 1, the pseudo-code (Supplementary Appendix S1), and our GitHub repository.

To explore the sensitivity and specificity of our resampling approach we simulated gene sets (see methods section 2) with varying GOI frequencies, and odds ratios for the two gene pairs (Supplementary Fig 2A). As expected, when we simulated two gene pairs with similar mutual exclusivities, no significant differences in their odds ratio distributions were detected (Fig 1C, Fig 1D left panel). Furthermore, large differences in simulated odds ratios required relatively few observations of a genetic event (Fig 1C, Fig 1D right panel). If the simulated gene pairs had a small difference in their simulated odds ratios, whether or not a strong difference was detected depended on the abundance of the GOI in the cohort (Fig 1D middle panel, Supplementary Fig 2B). For example, in a cohort of 500 patients, if the two positive control genes had an odds ratio of 0.05 and GOI and PC1 had an odds ratio of 0.2, at least 28 GOI patients were needed before a high difference in odds ratios could be detected, i.e. this cohort requires at least 28 GOI patients before we can conclude that the difference in odds ratios between the two gene pairs is real.

Our method is sensitive enough to detect meaningful pairwise differences in mutual exclusivity in ∼10/500 (2%) patients in a clinical cohort. Given that this sensitivity should be sufficient for the observed frequency of ALK^ATI^ (2-10%), we decided to re-examine the literature controversy surrounding this putatively oncogenic alteration.

### ALK^ATI^ is not as mutually exclusive as other established therapeutic targets in melanoma

The Couts paper contained a sample size of 6 ALK^ATI^ patients, but it identified NRAS and BRAF mutations in patients that harbored the ALK^ATI^ alteration (34). Thus, the lack of mutual exclusivity of ALK^ATI^ with the transforming melanoma oncogenes BRAF and NRAS has been suggested, but not conclusively and quantitatively demonstrated with an appropriately powered analysis. Pairwise comparison of conditional selection clearly showed that ALK^ATI^ is not as mutually exclusive with BRAF or NRAS as they are with each other (Fig 2A-C, see methods for description of data acquisition, sorting, and analysis). No significant difference in mutual exclusivity was detected when BRAF NRAS was compared to another known mutually exclusive gene pair (EGFR and KRAS in lung cancer, Fig 2D). Looking back at our simulated patient cohorts (black cross in Fig 1C-D), it is clear that this observed lack of mutual exclusivity of ALK^ATI^ is not confounded by its low abundance. If there were 12 ALK^ATI^ patients (not 38 ALK^ATI^ patients) in our dataset, we would have been unable to make any conclusions about whether a BRAF-NRAS level of mutual exclusivity exists in ALK^ATI^. Thus, ALK^ATI^ is significantly less mutually exclusive with BRAF or NRAS than they are with each other.

**Figure 2:**
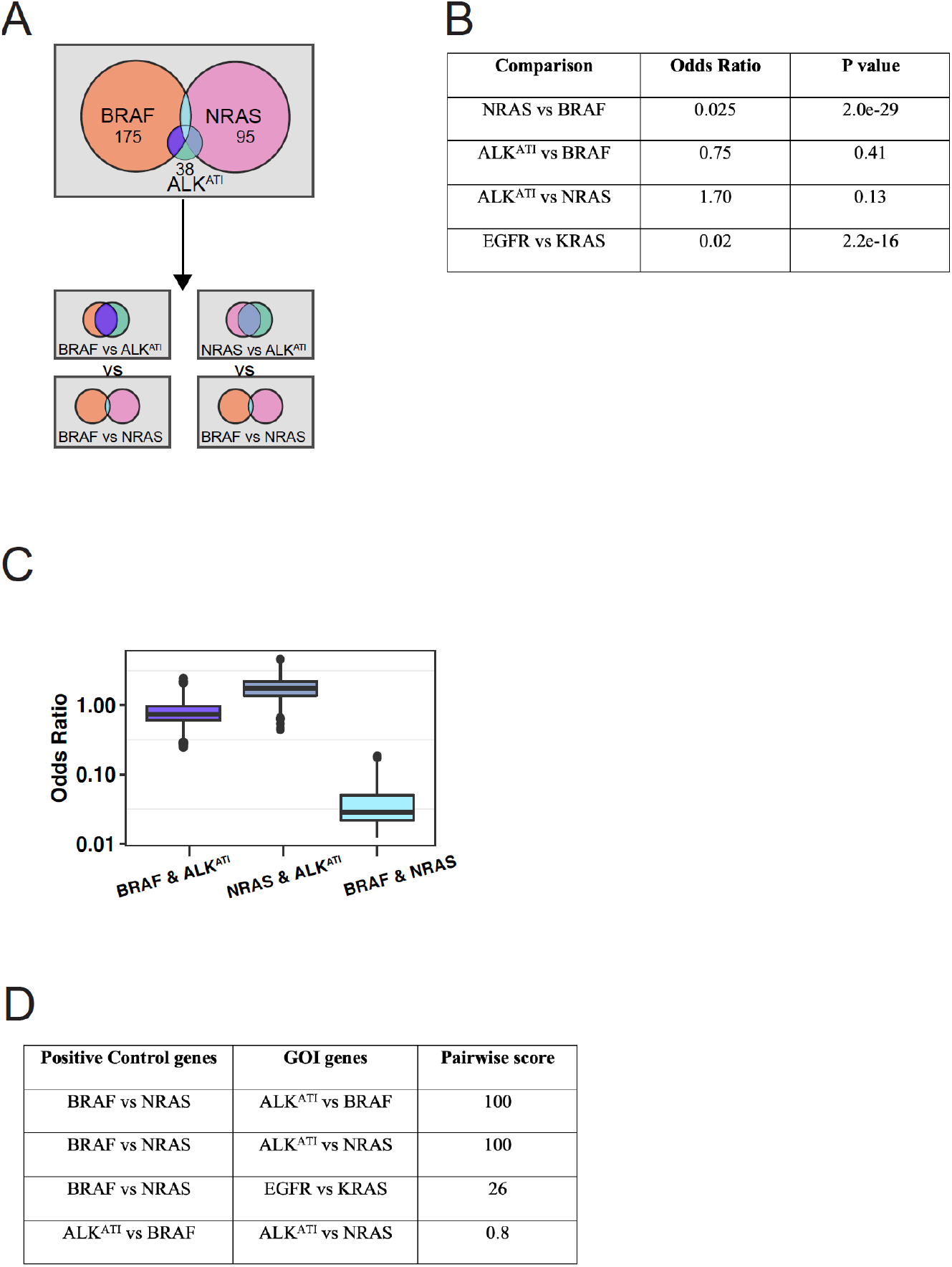
Performing pairwise comparisons of conditional selection on ALK^ATI^ reveals a lack of mutual exclusivity with transforming melanoma oncogenes BRAF and NRAS. **(A)** An illustration of the parameter space of ALK^ATI^ and the melanoma oncogenes BRAF and NRAS. The overall counts of the original data are: cohort size of 340 patients; abundances of ALK^ATI^, BRAF, NRAS: 38, 175, and 95. BRAF and ALK^ATI^: 12 patients, BRAF and NRAS: 25 patients, NRAS and ALK^ATI^: 8 patients. **(B)** ALK^ATI^ is not mutually exclusive with BRAF or NRAS. BRAF and NRAS in melanoma and EGFR and KRAS in lung cancer are mutually exclusive with each other (p-value from Fisher’s exact test). **(C)** Pairwise comparisons of the odds ratio distributions of ALK^ATI^ with BRAF, ALK^ATI^ with NRAS, and BRAF with NRAS. For BRAF and NRAS, the frequency of BRAF (51%) was reduced to match the frequency of ALK^ATI^ in the dataset (11%) according to our pairwise comparisons method. **(D)** Scores from the pairwise comparisons in **(C)**.

We also considered the possibility that the lack of mutual exclusivity of ALK^ATI^ with BRAF and NRAS was due to the definition of the initial filter cutoffs in the original RNA-seq analysis (the exact filters are mentioned in the methods). By systematically changing the cutoffs for all of the filters combinatorically, we varied the sensitivity for ALK^ATI^ detection by orders of magnitude (Fig 3A, top). More stringent filter sets resulted in fewer ALK^ATI^ calls, whereas less stringent filters resulted in more calls (Fig 3A, bottom). Performing pairwise comparisons of conditional selection on all combinations of these filter cutoffs never created mutual exclusivity. This filter analyses indicated that mutual exclusivity in AKL^ATI^ could not be observed with any data driven RNA-seq definition.

**Figure 3:**
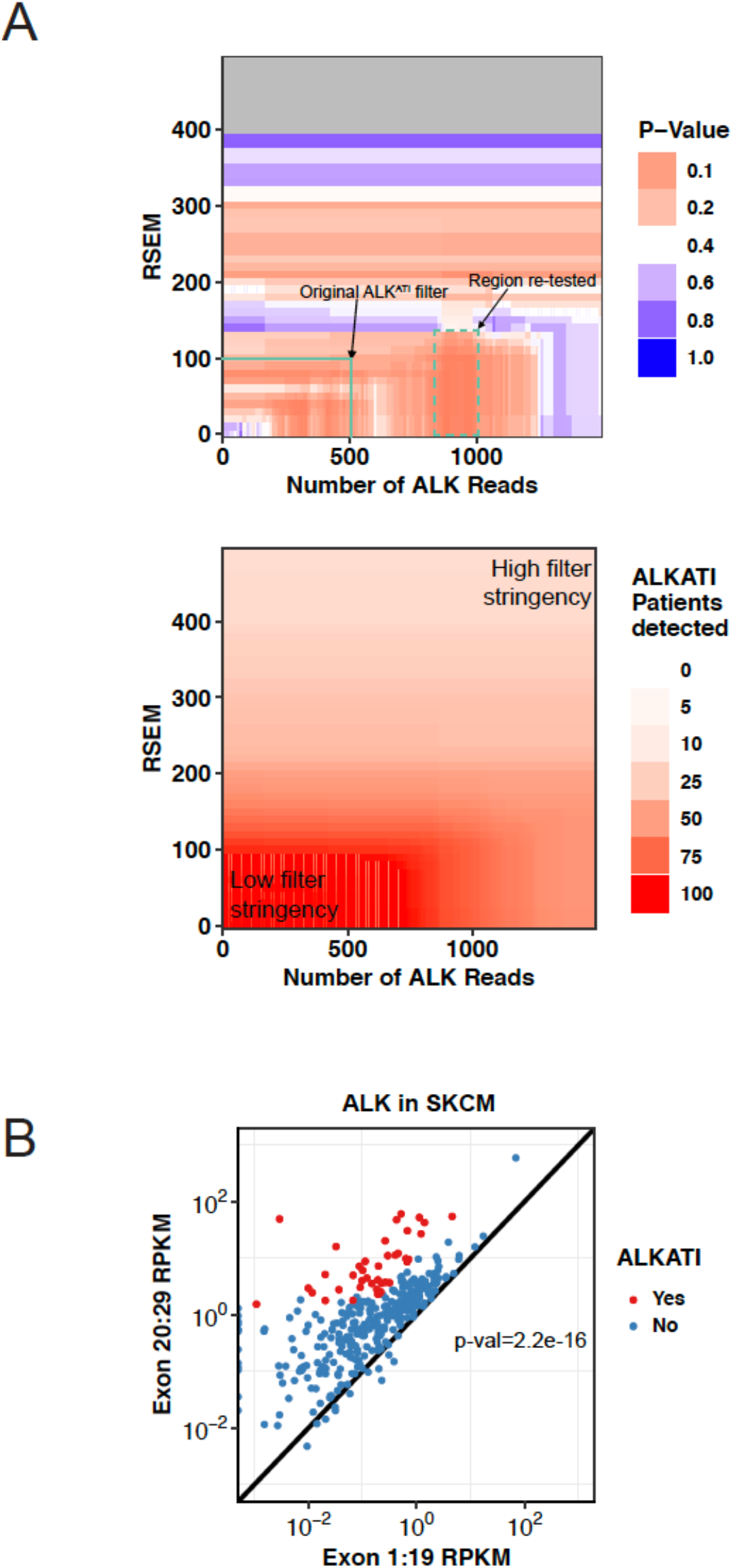
Changing the RNA-Seq filter cutoffs does not identify mutual exclusivity in ALK^ATI^. **(A)** Top: Changing the RNA-seq filter cutoffs for RSEM (43), RPKM (44), and count data did not result in mutually exclusive regions of ALK^ATI^ with BRAF or NRAS. Filters used were RSEM 10-1000, counts 100-1000, ex20-29/ex1-19 ratio 10-100. Only the minimum p-value observed amongst all exon ratios tested is displayed. The midpoint of the p-value color gradient is 0.3. Regions that filtered for <10 ALK^ATI^ patients were not included in the pairwise comparison analysis and are colored gray. Bottom: The number of patients that are categorized as ALK^ATI^ positive decreases as the filter stringency increases. **(B)** The kinase domain of ALK (ex. 20-29) is significantly overexpressed in the majority of SKCM patients (p-value is from a χ^2^ test).

This analysis shows the utility of tests for pairwise conditional selection to demonstrate quantitative differences in the degree of mutual exclusivity between a positive control gene pair and a new and potentially oncogenic alteration. We also strongly demonstrate a lack of mutual exclusivity between ALK^ATI^ and established MAPK pathway driver mutations.

### Kinase domain expression imbalance in ALK is nearly ubiquitous in melanoma and lung cancers

Given the conclusive lack of mutual exclusivity between ALK^ATI^ and transforming melanoma oncogenes, we decided to look deeper at the initial signal, i.e. the bias towards ALK expression in the kinase domain (Exons 20:29). We expected to see a distribution centered at equivalent expression between the kinase domain and the upstream coding region (the diagonal line in Fig 3B). We posited that expression levels in exons 1-19 and 20-29 should be distributed above and below the diagonal, with ALK^ATI^ patients being strong outliers from this expected relationship. However, we observed a significant bias towards overexpression of the kinase domain of ALK across all melanoma patients, p-value from Kolmogorov-Smirnov test: 2.2e-16. In fact, almost all the skin cutaneous melanoma (SKCM, n=340) and lung adenocarcinoma (LUAD, n=477) patients in the TCGA (Fig 3B, Supplementary Fig 3A) expressed higher levels of the ALK kinase domain than exons 1-19. The ubiquity of this deviation in all patients led us to suspect that a systematic error could be at play. While multiple interpretations of this signal exist, it is concerning that all patients have some propensity to overexpress the kinase domain. As a control, we also examined whether an imbalance of expression towards the kinase domain is a unique feature of ALK. We did not see this propensity for kinase domain expression in EGFR in SKCM or in EGFR in LUAD, p-value from KS-test:0.66 (Supplementary Fig 3B). Hence, the kinase domain expression imbalance is not a generalizable finding and raises concerns of potential systemic biases in exon specific expression levels in ALK in the TCGA.

### ALK^ATI^ is not sufficient for growth/transformation *in vitro*

Our computational re-analysis of ALK expression data in melanoma suggested that ALK^ATI^ is significantly less mutually exclusive with NRAS and BRAF than they are with each other. In the original Wiesner *et al*. study (32), ALK^ATI^ was argued to be sufficient for transformation of Ba/F3s to growth factor independence. When we transduced Ba/F3s with ALK^ATI^ and EML4-ALK (9 independent replicates each), we found that ALK^ATI^ took significantly longer to grow out (two population doublings in 6.7±0.4 days for EML4-ALK and 10.0±1.0 days for ALK^ATI^, Fig 4A). However, we also reasoned that longer outgrowth times were indicative of a weaker transforming potential. As such, we scaled up the number of transductions to 48 independent replicates by performing many parallel transductions of ALK^ATI^, EML4-ALK, and vector. The results were striking. Growth factor independence was observed in 100% of EML4-ALK replicate transductions, but only 37.5% ALK^ATI^ replicate transductions, and in 16% of vector controls (Fig 4B). Viral titers were essentially indistinguishable across these three constructs (Supplementary Fig 4A); puromycin resistance cassette transfer as indicated by time to outgrowth following puromycin selection was similar as well. Thus, ALK^ATI^ exhibits only a modest increase in transformation efficiency when large scale replication is performed.

**Figure 4:**
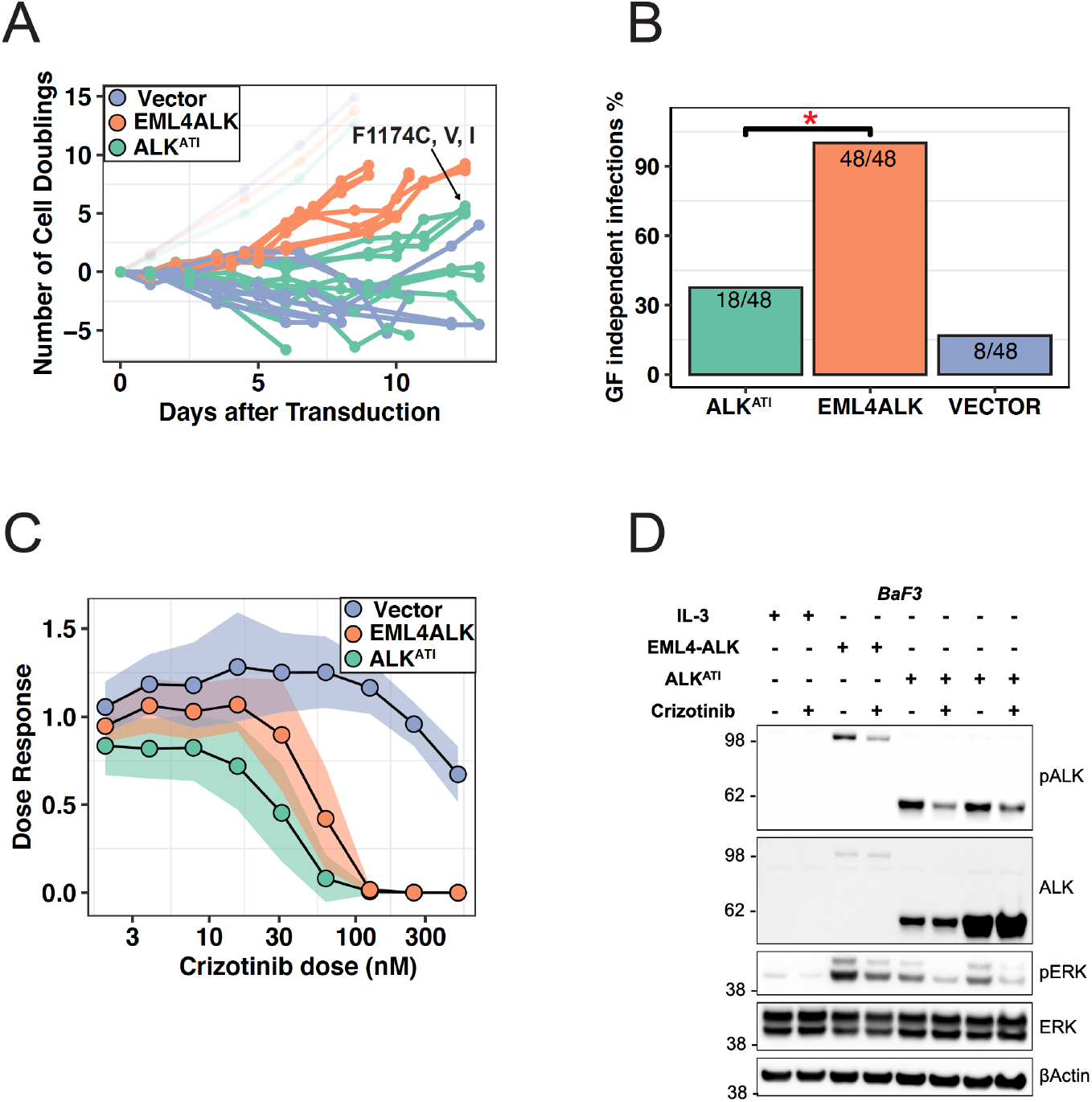
ALK^ATI^ is not sufficient in melanoma and does not predict therapeutic dependency. **(A)** Time to growth factor independence, as measured by number of cell doublings, between ALK^ATI^-transduced Ba/F3 cells (green), EML4ALK-transduced cells (orange), and vector-transduced cells (purple). Faded lines represent transduced cells growing without selection (grown with IL3). Activating mutations in the kinase domain of ALK were detected after sequencing the genomic DNA of all growth factor independent outgrowths (see Supp. Fig. 4B). **(B)** Proportion of infections that transformed for growth factor independence (χ^2^ test p-value for ALK^ATI^ vs EML4ALK: 2.2e-10, ALK^ATI^ vs Vector: .038). **(C)** Crizotinib dose-response study for IL3 independent EML4-ALK, ALK^ATI^, and IL3 dependent vector-transduced Ba/F3 cells. Data are mean ± 95% confidence intervals (shaded region). 3, 5, and 1 independent transductions for EML4-ALK, ALK^ATI^, and vector control were tested. For all cell lines, 3 technical replicates were used at each concentration, and each assay was repeated on 3 separate days. **(D)** Immunoblots showing ALK-overexpression and crizotinib dependence of EML4-ALK, and ALK^ATI^ Ba/F3 cell lines (n=2 independent transductions each). The bands represent EML4ALK (117kDa), ALK^ATI^ (67kDa), β-actin (42), and ERK1/2 (42,44 kDa). Immunoblots were repeated with three times with similar results. 1ug of lysate was loaded per well. An exposure time of 120s was used.

Next, we reasoned that growth factor independence in ALK^ATI^ Ba/F3 cells was infrequent because it required a relatively rare second genetic event. To test this hypothesis, we extracted the genomic DNA of the 18 ALK^ATI^ Ba/F3 cell lines that achieved growth factor independence and sequenced their ALK kinase domain. Surprisingly, 3 of the 18 ALK^ATI^ cell lines had well known transforming mutations in the ALK kinase (F1174C, F1174V, and F1174I, Fig 4A, Supplementary Fig 4B). These point mutations are the primary cancer causing mutations in ALK-mutated neuroblastoma and have been shown to constitutively activate the ALK kinase (36). These mutations are also sensitive to some ALK kinase inhibitors. This gives a concrete rationale for why ALK^ATI^ could transform Ba/F3 cells, and that those cells were sensitive ALK inhibition (32). We hypothesize that these mutations spontaneously arise in a small subset of transduced Ba/F3 cells, but that IL3 withdrawal strongly selects for the growth benefit conferred by these artifactual mutations. While the mutations were not found in all cells, their existence in 3 independent selections strongly suggests that the transformation results and therapeutic treatments of ALK^ATI^ can be confounded by secondary genetic events. To confirm that these mutations in ALK^ATI^ are sufficient for transformation, we added the 3 mutations to the original ALK^ATI^ plasmid via site directed mutagenesis. When we performed new transformation experiments, ALK^ATI-F1174 C, V, and I^ mutant cells transformed Ba/F3’s in a highly efficient manner that resembled our EML4-ALK positive control (Supplementary Fig 4C). To further confirm that these cells were ALK addicted, we treated Ba/F3 cells with crizotinib and examined their dose response curves and signaling state via western blots. ALK^ATI^ transformed Ba/F3 cells are sensitive to crizotinib and brigatinib (Fig 4C, Supplementary Fig 4D). Crizotinib treatment decreases phospho-ALK and phospho-ERK in a manner similar to EML4-ALK (Figure 4D). Together, this data suggests that while ALK^ATI^ transformed Ba/F3 are sensitive to crizotinib, they require a second transforming event (i.e F1174 mutations) to fully transform cells *in vitro*. These mutations are not seen in melanoma patients. Alongside our re-analysis of the genomic data and the Couts *et al* data (34), our transformation data strongly suggested that ALK^ATI^ is not sufficient for growth/transformation *in vitro*, and that ALK inhibitors will not be a good single agent treatment strategy in ALK^ATI^ positive melanoma. Though other transformation experiments were performed in (34), we argue that the large scale replication of the Ba/F3 results in our lab cast significant concerns on the other transformation studies in ALK^ATI^.

### ALK^ATI^ cannot rescue melanoma cell lines

While our Ba/F3 analysis suggests that the ALK^ATI^ alterations are not sufficient to transform tool cell lines to growth factor independence, we decided to test the potential oncogenicity of ALK^ATI^ in a more realistic melanoma cell line model. We transduced ALK^ATI^ into two BRAF_V600E_-harboring melanoma cell lines, and challenged them with a BRAF_V600E_ inhibitor, vemurafenib. The experimental rationale was simple, BRAF_V600E_ is necessary for melanoma cell survival, and sufficient for transformation (21,31). The drug vemurafenib inhibits only the mutant BRAF V600E protein as a monomer (16,31,37), while receptor tyrosine kinases signal through dimeric RAF family proteins (38). A candidate oncogenic RTK in melanoma with strong transforming potential should be able to rescue BRAF_V600E_ melanoma from a BRAF_V600E_ inhibitor.

We transduced EML4-ALK, ALK^ATI^, and vector into two different skin cancer cell lines, SKMEL28 and G361, both of which have a transforming V600E point mutation and are sensitive to vemurafenib (35). When we did this, ALK^ATI^ and vector-transduced melanoma cells had statistically indistinguishable vemurafenib dose responses (p-value: 0.49 for SKMEL28 and 0.97 for G361, Fig 5A). Western blots confirmed ALK^ATI^ constructs robustly expressed, and that both melanoma cell lines exhibited strong overexpression of the ALK^ATI^ construct (Figure 5B-C). The inability of ALK^ATI^ to rescue melanoma cell lines is in line with the notion that ALK^ATI^ is not an oncogenic driver in melanoma.

**Figure 5:**
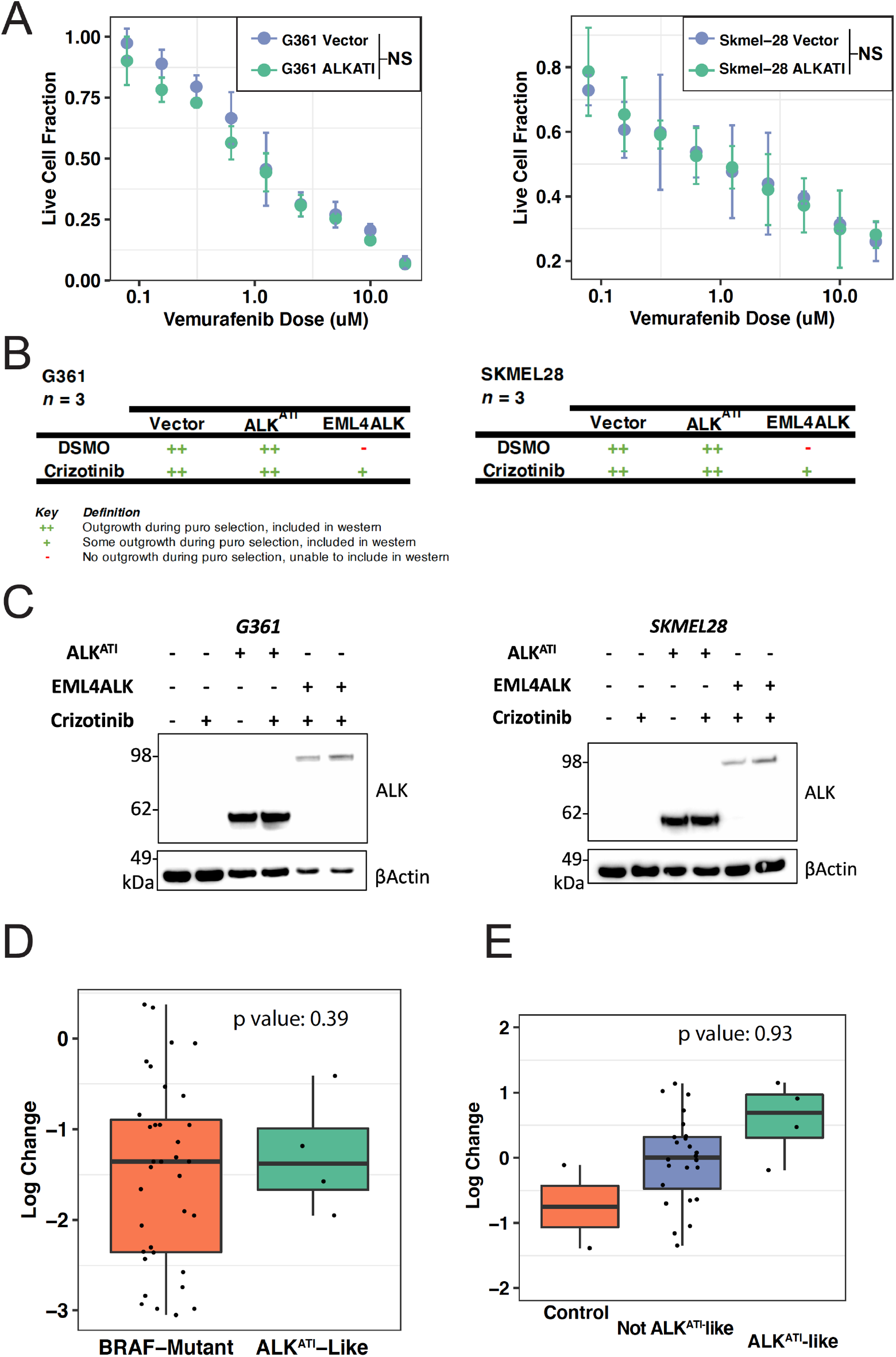
ALK^ATI^ cannot replace known oncogenes in melanoma. **(A)** Dose response of ALK^ATI^ and vector-transduced SKMEL-28 and G-361 cell lines. ALK^ATI^ does not improve the dose response of SKMEL-28 (p-val: 0.49) and G361 (p-val: 0.99) to vemurafenib. P-value calculated using a one-sided paired t-test test between ALK^ATI^ and vector. Error bars represent standard deviation on 3 replicates of 3 independent transductions (9 total replicates per condition per concentration). **(B)** Outcome of stable overexpression of Vector control, ALK^ATI^, and EML4-ALK on SKMEL-28 and G361 cell lines. 3 independent transductions were tested for each condition. **(C)** Immunoblots for ALK^ATI^ overexpression in melanoma. Two melanoma cell lines transduced with vector control, ALK^ATI^, or EML4-ALK. Two independent tranductants are shown for EML4-ALK and ALK^ATI^. The bands represent EML4ALK (117kDa), ALK^ATI^ (67kDa), and β-actin (42). Immunoblots were repeated with three times with similar results. 10ug of SKMEL28/G361 lysate were loaded. A crizotinib dose of 100nM was used. An exposure time of 120s was used. **(D)** Comparing the vemurafenib dose response of BRAF-mutated cell lines (orange, n=43), and BRAF-mutated cell lines that have an ALK^ATI^-like expression (cyan, n=4). P-value is from a one-sided multiple linear regression of vemurafenib intolerance mapped as a function of ALK RSEM, RPKM, and Exon expression. **(E)** Crizotinib sensitivity of melanoma cell lines with ALK^ATI^-like expression (cyan, n=4) or not (purple, n=32). Crizotinib sensitive control cell lines (orange) are Kelly (ALK^F1174L^ neuroblastoma), and NCI-H228 (EML4-ALK in lung adenocarcinoma). P-value is from a one-sided multiple linear regression of crizotinib sensitivity mapped as a function of ALK RSEM, RPKM, and Exon expression.

We were unable to transduce SKMEL28 and G361 cells with EM4-ALK on multiple transduction attempts. However, we decided to proceed with our vemurafenib challenge in the absence of EML4-ALK. EML4-ALK is an established oncogenic drivers in NSCLC and melanoma (12,39). We confirmed our viral titer and infectivity by simultaneously transducing and transforming Ba/F3 to IL-3 independence, as well as infecting Hek293T and selecting for puromycin resistance. Interestingly, virus packaged with the EML4-ALK oncogene readily transformed Ba/F3 cells, easily infected Hek293T cells, and selected with puromycin, but we could not successfully select for EML4-ALK containing SKMEL28 or G361 cells (Supplementary Table 1). Because this was a negative result, we tested the idea that conditional selection via mutual exclusivity might be occurring. Our hypothesis is that simultaneous expression of two growth pathway activating variants in the same cell is not well tolerated (28,29). To test this, we performed the same infections that are described above, but during the selection of the infected cells, we performed all selections in both the presence and absence of crizotinib. Whereas vector control and ALK^ATI^ cells formed stable cell lines in the presence and absence of crizotinib, we were only able to stably select melanoma cells expressing EML4-ALK when we selected for transductants in the presence of the ALK inhibitor crizotinib at 100nM (Figure 5B, Supplementary Table 1). This suggests that melanoma cells require ALK inhibition in order to stably express EML4-ALK. The crizotinib dependence of the known oncogene EML4-ALK and the crizotinib independence of ALK^ATI^ in melanoma cells alongside the inability of ALK^ATI^ to rescue vemurafenib challenge suggest that ALK^ATI^ does not act as a constitutively active oncogene in melanoma cells. It also suggests that the expression of an activated ALK construct and BRAF_V600E_ actually inhibits melanoma growth.

### ALK expression imbalance does not predict transforming potential or single agent therapeutic dependency

We further probed the transforming potential of ALK^ATI^ by searching for evidence for an ALK^ATI^ dependent transforming potential *in vitro*. To do this, we analyzed BRAF-mutant cell lines (n=43) from CCLE (35) for their ALK expression and dose response to BRAF-inhibitors. In this dataset, 4 of these 43 cell lines were ALK^ATI^-like (exceeded 2/3 filters). None of these 4 cell lines and a substantial difference in their sensitivity to 11 distinct BRAF inhibitors (Fig 5D, Supplementary Fig 5A). Furthermore, the degree of exon imbalance in ALK expression (when treated as a continuous variable) did not predict improved survival against a BRAF-inhibitor challenge.

While previous experiments suggested that ALK^ATI^ may not be sufficient for oncogenesis, we still wondered if ALK^ATI^ conferred sensitivity to ALK inhibitors in melanoma cell lines *in vitro*. To test this, we analyzed melanoma cell lines (n=32) from CCLE (35), 4 of which were ALK^ATI^-like (exceeded 2/3 filters). No evidence of ALK-inhibitor sensitivity was found in any of the 4-ALK^ATI^-like cell lines (Fig 5E, Supplementary Figure 5B). Furthermore, the degree of exon imbalance in ALK expression did not predict ALK-inhibitor responses in these 32 cell lines (linear regression p-value for crizotinib: 0.93). Hence, the degree of observed exon imbalance is not correlated with single agent sensitivity to ALK-inhibition across 32 melanoma cell lines.

## Discussion

Conditional selection has been used to study mechanisms of oncogenic activation in a variety of contexts. Previous research has used mutual exclusivity as an indicator of oncogenic network modules and dysregulated pathway analyses, and to identify evolutionary dependencies from alteration occurrences in pan-cancer analyses. Importantly, mutual exclusivity is often used to prioritize rare genomic findings without defining statistical power to detect a given effect size, or to compare to an expected effect size. To our knowledge, no previous method accurately quantifies a negative signal for conditional selection in rare variants. Our code base allows us to quantify a negative finding relative to how often a lack of mutual exclusivity would be seen in a positive control gene pair. Our simulations show that our method for pairwise comparisons is a quantitative and statistically robust method to identify a lack of conditional selection with sufficient statistical power.

We applied our pairwise comparisons of conditional selection to a transcript alteration in ALK^ATI^ because of the controversy in the literature (32,34). Our analysis clearly demonstrates a lack of conditional selection for ALK^ATI^. ALK^ATI^ is significantly less mutually exclusive than BRAF and NRAS are with each other in melanoma. Moreover, our experiments suggest that ALK^ATI^ is not sufficient for cellular transformation, that kinase domain imbalance does not predict inhibitor response, and that single agent ALK inhibition is unlikely to be therapeutic in melanoma cells.

In their original paper, Wiesner et. al. (32) performed a substantive and detailed description of the ALK^ATI^ event. They found enrichment of H3K4me3 and RNA Pol II near the ATI transcription initiation site of tumors expressing ALK^ATI^ (32). Wiesner *et al* also confirmed that ALK^ATI^ is expressed at both ALK alleles by comparing DNA, RNA, and H3k4me3 levels. They also performed gene expression profiling of RNA-Seq datasets showing that ALK^ATI^-like expression is found in 2-11% of melanoma samples and sporadically in various other tumor types and not in normal tissue samples (32,33). The breadth and depth of the analysis leads us to believe that ALK^ATI^ is likely a true transcript variant. Moreover, while we believe our work (alongside the work of Couts *et al* (34)) provides strong evidence against single agent therapy, it would be unfair to ignore the potential for therapeutic relevance of ALK^ATI^ in contexts other than ALK inhibitor monotherapy. Moreover, from a biological perspective, it would also be unfair to rule out some sort of unknown biological role for ALK^ATI^ that does not fit conventional definitions of driver oncogenes. However, combining Couts’ PDX data with the questionable transformation potential of ALK^ATI^ (34), the lack of objective responses in ALK^ATI^ expressing melanomas to ALK inhibitors, and the clear the lack of mutual exclusivity of ALK^ATI^ in melanoma casts significant doubt upon the single agent therapeutic rationale for ALK^ATI^.

In the melanoma landscape, dramatic responses to approved and investigational immunotherapy agents are yielding important steps forward for patient care. We have systematically shown that the original ALK^ATI^ data should be re-evaluated in light of our data reproducing the original finding, and in light of the compelling recent reports in PDX models (34,40). Combined with the fact that the patient data in the original manuscript did not achieve the typical clinical criteria for an objective partial response, we strongly recommend that single agent ALK inhibitors receive no further testing in ALK^ATI^ patients when other investigational or off label options exist. Given the weight of evidence, it seems unlikely that refractory patients will benefit from this treatment. We also suggest that pairwise tests for conditional selection will be a useful tool to triage any rare genomic finding in the mountains of cancer sequencing data generated in late-stage patients.

## Methods

### 1.1 Frequency correction in gene pairs

Suppose we have a joint table from our positive controls with frequencies

The odds ratio, defined as the strength of association between two events, is (*p*_11_ * *p*_00_)/(*p*_10_ * *p*_01_) for Table 1 above.

**Table 1:**
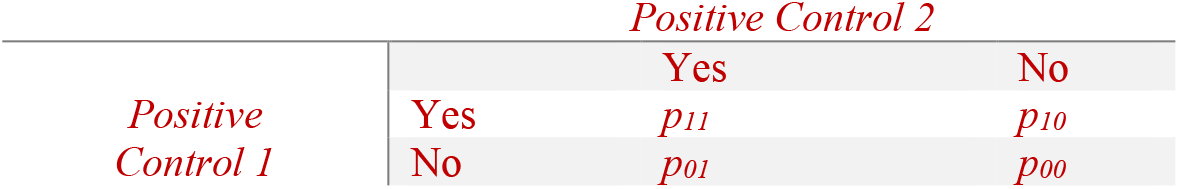
A 2×2 Contingency table of the positive control 1 and positive control 2 genes. Where *p*_*11*_ refers to the frequency of the population having mutations in both the positive control 1 gene and the positive control 2 gene; *p*_*10*_ refers to the population having a mutation in the positive control 1 gene only, and so on.

Also suppose we have a GOI with frequency *p*_*GOI*_. We would like to transform the above table to a new table such that *Positive Control 1* positive individuals appear at frequency *p*_*GOI*_, but where the odds ratio remains the same. We can do this by adding or removing the appropriate fraction of *Positive Control 1* positive individuals without regard to their status with respect to *Positive Control 2*. This results in a new table with cell probabilities given by:

This table is derived by multiplying each row in the original table by an appropriate constant (e.g., the first row is multiplied by the ratio between the desired fraction of *Positive Control 1* positive individuals *p*_*GOI*_ and the observed fraction in the *p*_10_+*p*_11_ from the original table). It is easy to verify that the table 2 is a valid probability distribution with cells that sum to 1, that *Positive Control 1* is now at frequency *p*_*GOI*_, and that the odds ratio is unchanged from the original table.

**Table 2:**
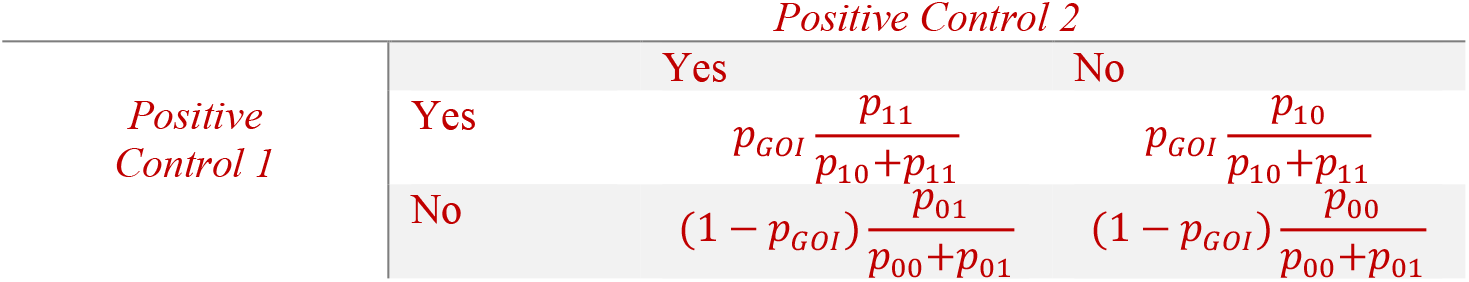
A 2×2 Contingency table showing of the positive control 1 and positive control 2 genes with the frequency of the positive control 1 gene corrected to the frequency of the gene of interest.

### 1.2 Pairwise comparisons of gene pairs

In order to estimate the typical range of odds ratios likely to be produced under the hypothesis that GOI and PC2 are as mutually exclusive as PC1 and PC2 while controlling for the observed frequency of GOI, we can thus construct samples from the above table of size *N*, where *N* is the size of the cohort of our GOI by PC2. Specifically, we construct these samples as a draw from a multinomial distribution with *N* trials and probabilities given by the above table of probabilities where the expected frequency of *Positive Control 1* positive individuals has been set to *p*_*GOI*_. Independently drawing 1000 such tables and calculating the odds ratio for each table, we calculate the percent of PC1 vs PC2 odds ratios that have a lower odds ratio than the odds ratio for PC1 and GOI (labeled as score in Fig 1). The higher this score, the lower the overlap is between the odds ratios of the two gene pairs. A score of >95% was set as the threshold for rejecting the hypothesis that GOI and PC1 are as mutually exclusive as PC1 and PC2. Section 1 of the pseudo-code contains a step-by-step implementation of this process.

#### 2. Generating simulated cohorts at various mutual exclusivities and abundances

We simulated cohorts of gene pairs to test our pairwise comparisons method and to characterize its sensitivity. Suppose we would like to simulate a joint table from our positive controls with frequencies

Also suppose we would like to simulate a joint table with the gene of interest and positive control 1 gene with frequencies

Here, we outline a method to calculate the frequencies in each quadrant of contingency tables 3 and 4 using three inputs:

**Table 3:**
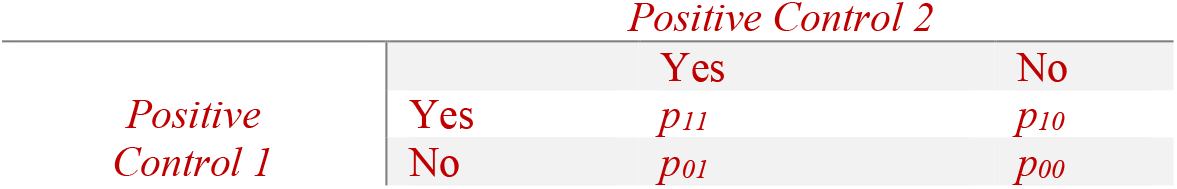
A 2×2 Contingency table of the positive control 1 and positive control 2 genes. Where *p*_*11*_ refers to the frequency of the population having mutations in both the positive control 1 gene and the positive control 2 gene; *p*_*10*_ refers to the population having a mutation in the positive control 1 gene only, and so on.

**Table 4:** A 2×2 Contingency table of the positive control 1 gene and the gene of interest. Where *q*_*11*_ refers to the frequency of the population having mutations in both the positive control 1 gene and the gene of interest; *q*_*10*_ refers to the population having a mutation in the positive control 1 gene only, and so on.

|  |  | *Gene of interest* |  |
| --- | --- | --- | --- |
|  |  | Yes | No |
| *Positive Control 1* | Yes | $q_{11}$ | $q_{10}$ |
| | No | $q_{01}$ | $q_{00}$ |

**Table 5:**
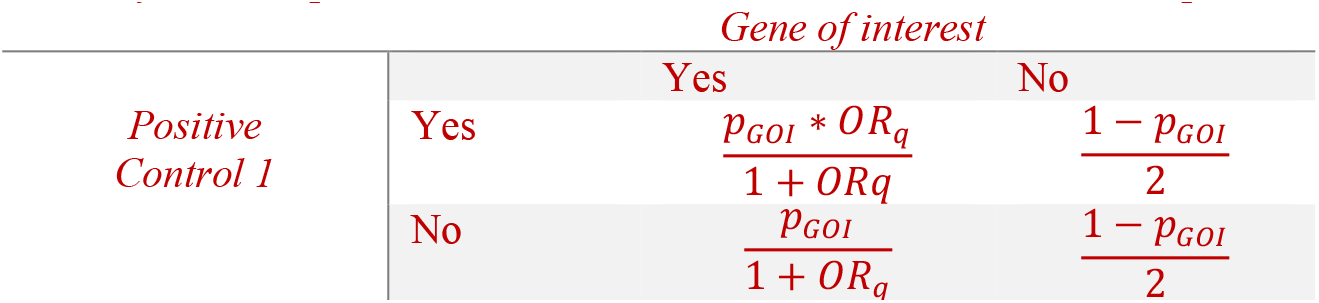
A 2×2 contingency table of the positive control 1 gene and the gene of interest in which each frequency is a function of the input variables *OR*_*q*_ and *p*_*GOI*_.

**Table 6:**
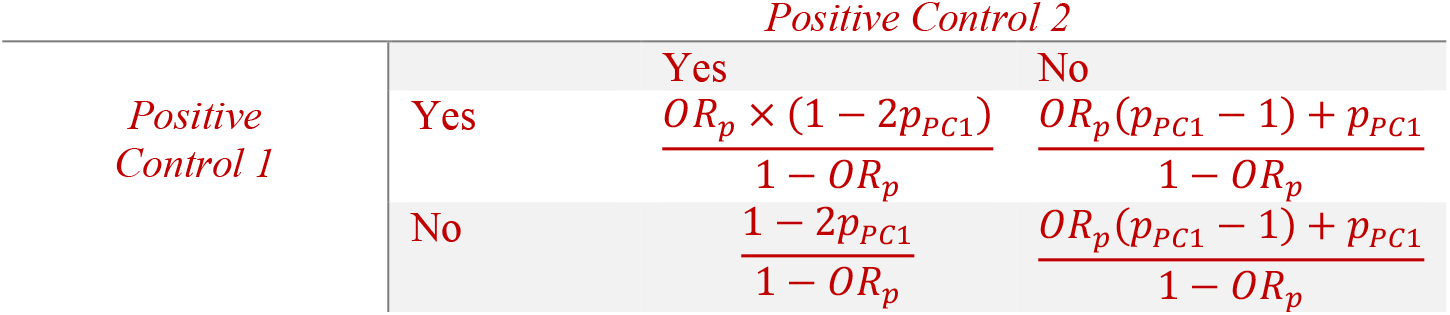
A 2×2 contingency table of the two positive control 1 genes in which each frequency is a function of the input variables *OR*_*p*_ and *p*_*PC1*_.

- The odds ratio of the positive control genes, *OR*_*p*_, which can be written in terms of the frequencies in table 3:

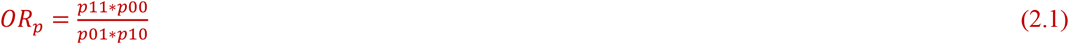
- The odds ratio of the gene of interest and positive control 1, *ORq*, which can be written in terms of the frequencies in table 4:

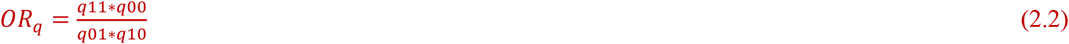
- The frequency of the gene of interest, *p*_*GOI*_, which is the sum of its two compartments in table 4:

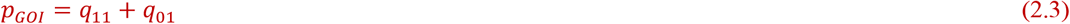

Since we are simulating valid probability distributions, the frequencies in table 3 and 4 should sum to 1. Hence,

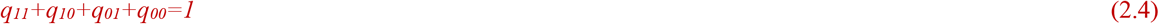

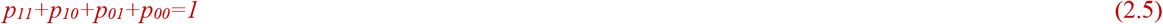

For simplicity, we assume that the frequency of patients without a mutation in either gene is the same as the frequency of patients with a mutation in positive control 1 only:

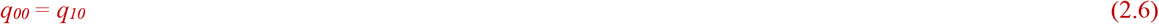

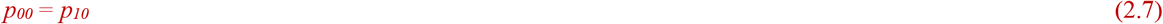

Eq 2.6 can be substituted into eq 2.2:

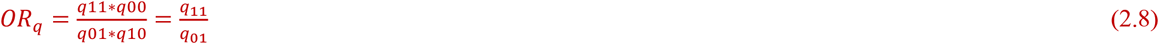

Similarly, Eq 2.7 can be substituted into eq 2.1:

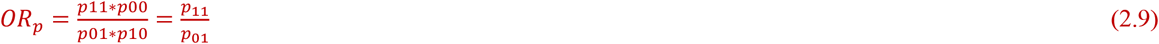

Eq 2.3, 2.4, 2.6 and 2.8 can be arranged as a system of linear equations and solved using Gaussian elimination:

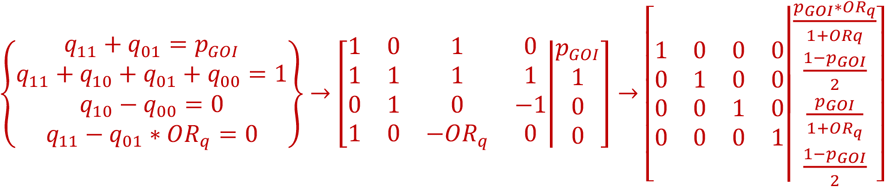

This way, the compartments of table 4 are solved as functions of the inputs:

Since assume that both joint tables come from the same patient cohort, the overall frequency of positive control 1 gene, termed *p*_*PC1*_, is the same in both Tables 2 and 3. Therefore, *p*_*PC1*_ can be rewritten as a function of the inputs *p*_*GOI*_ and *OR*_*q*_:

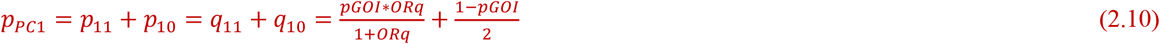

Therefore, eq 2.10, 2.5, 2.7, and 2.9 can be arranged as a system of linear equations and solved using Gaussian elimination:

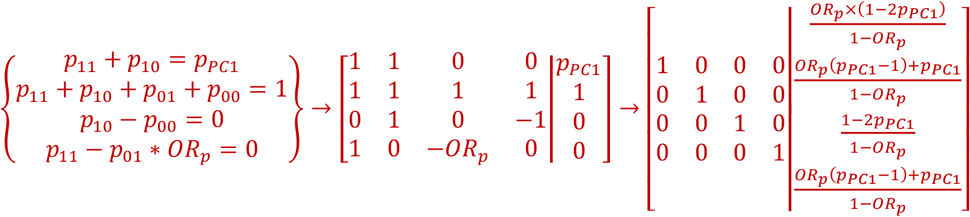

This way, the compartments of table 3 are solved:

Having established a method to calculate frequencies in contingency tables 3 and 4 using the three inputs *OR*_*p*_ *OR*_*q*_ and *p*_*GOI*_, next, we simulated these contingency tables at various inputs. Each set of these input variables generates a contingency table for the positive control 1 gene and the positive control 2 gene, and another contingency table for the positive control 1 gene and the gene of interest. The odds ratio of the positive control, genes *OR*_*p*_, was varied from 0.01 (very mutually exclusive) to 0.1 (less mutually exclusive). The odds ratio of the gene of interest gene with the positive control 1 gene, *OR*_*q*_, was varied from 0.01 (mutually exclusive) to 1 (not mutually exclusive). The frequency of the gene of interest, *p*_*GOI*_, in the cohort was varied from 0.005 (rare) to 0.5 (abundant). Supplementary Fig 2A provides a visual illustration of how these input variables affect the simulated cohorts. Once these simulated contingency tables of gene pairs were generated, the previously described pairwise comparisons method (methods section 1) was used to determine whether GOI is as mutually exclusive with PC1 as PC1 and PC2 were with each other. Section 2 of the pseudo-code contains a step-by-step implementation of this process.

#### 3. Analysis of public data sets

We downloaded our level 3 TCGA data from the Broad Institute TCGA GDAC Firehose (http://gdac.broadinstitute.org/) 2016_01_28 run. In our original ALK expression filters (Fig 2), ALK^ATI^ candidates were identified as samples with an ALK expression level of RSEM ≥ 100, ≥ 500 total reads across all ALK exons, and ≥ 10-fold greater average expression in exons 20–29 compared to exons 1–19. When varying filters, ALK^ATI^ patients were identified as samples with an ALK expression level of RSEM≥100, number of ALK reads≥500, and Exon20-29/Exon1-19 RPKM ratio≥10 (Fig 3). Cell line expression and drug sensitivity data (Fig 4) was downloaded from the cancer cell line encyclopedia (https://portals.broadinstitute.org/ccle/data). Melanoma cell-lines were classified as ALK^ATI^-like if they matched 2/3 ALK^ATI^ filters.

Cell line expression and drug sensitivity data: CCLE 2019 RNA-seq gene expression data for 1019 cell lines were downloaded from the Broad Institute CCLE website ((https://portals.broadinstitute.org/ccle). We used this data to identify ALK^ATI^-like cell lines for their sensitivity to BRAF and ALK-inhibitors in melanoma. For 49 melanoma cell lines, ALK expression data was extracted and ALK^ATI^ expression was detected using two criteria: 1) ALK is expressed in the cell line with RSEM> 100, and read counts > 500, and 2) Since the ATI-site resides in intron 19 of ALK, a 10-fold greater expression of ALK exon20-29 than exon1-19 was expected for ALK^ATI^-like cell lines.

For the identification of EML4-ALK cell lines, CCLE 2019 Fusion calls for 1019 cell lines were downloaded from CCLE. The EML4-ALK fusion calls was identified in 2 lung cancer cell lines(NCIH3122 and NCIH2228) and a pancreas cancer cell line(SNU324). For the identification of BRAF-mutant cell lines: 2019 cancer cell line mutant calls were downloaded from CCLE. A total of 111 BRAF mutants cell lines were identified that were targeted by 11 BRAF inhibitors.

#### 4. *In vitro* transformation and drug treatment assays

We used a standard Ba/F3 transformation protocol for lentiviral transduction. Lentiviral particles were made by transfecting HEK293T cells with the plasmid of interest and with packaging plasmids (3^rd^ generation Lentiviral system from Addgene). Replication incompetent virus was collected using a BL2+ safety protocol. Upon virus collection, Ba/F3 cells at 500k/mL in 4mLs were infected with an equal volume of virus. A minimum of three replicates were used for each infection condition. Three days after infection, the Ba/F3s cells were spun out of virus and selected for IL3 independence and/or puromycin resistance (0.5ug/mL Puromycin was used). An infection was determined to be successful Ba/F3 cells for a given construct grew out for both IL3 independence and puromycin resistance. During selection, cell growth was quantified by doing daily counts of live/dead cells on a daily basis. These live/dead analyses were performed by supplementing 20uLs of cell culture with 0.4% trypan blue and subsequently counting live cells using a hemocytometer. Alternatively, live dead cells were counted using flow cytometry analysis (BD Accuri C6 Plus).

#### 5. Lentiviral Particle Quantification

For each infection, before transducing Ba/F3s with virus, 500uL of virus was set aside and frozen at -20C. The number of lentiviral particles were quantified using the QuikTiter Lentivirus Quantitation Kit from Cell Biolabs (HIV P24 ELISA). All reported viral titers were within the linear range of the standard curve made using a positive control. Transduction efficiency was verified by counting outgrowth rates using puromycin selections across multiple infections.

#### 6. Cell Lines

Cell lines used were: SK-MEL-28 (ATCC HTB-72), G361 (ATCC CRL-1424), Ba/F3 (DSMZ ACC-300), HEK-293T (ATCC CRL-1573). Prior to use, all cell lines were tested to be free of mycoplasma using a biochemical-based test (Mycoalert plus, Lonza). Ba/F3 cell lines were cultures in RPMI while HEK-293T, Skmel-28 and G361 cell lines were cultured in DMEM. Media was supplemented with a final concentration of 10% FBS, and 1% Penicillin/Streptomycin/L-Glutamine. Wt Ba/F3 cell lines were grown in media supplemented with 1ng/mL IL3 (R&D systems). Stable transductions were verified for puromycin resistance by using kill curve concentrations ranging from 0.25ug/mL to 2ug/mL.

Cells were split when they were 70-80% confluent. The subculture ratios used for all of our cell lines included splitting ratios of 1:5 every 2-4 days when cells were 70-80% confluent.

Ba/F3_Wt_ cell lines were grown in 10ng/mL IL3. After transduction with an oncogene, cell lines were selected with Puromycin to test transduction efficiency and subsequently with -IL3 to assess growth factor independence.

#### 7. Plasmids

The sequence of ALK^ATI^ was obtained from the European Nucleotide Archive under the expression number LN964494. The sequence of EML4-ALK was obtained from Genbank AB274722.1. The coding sequences, cloned into a pLVX-IRES-Puro backbone, were prepared by Genscript Gene Services. Please refer to Supplementary Table 2 for the sequence of both of these plasmids.

#### 8. Immunoblots

Cells were lysed in RIPA buffer (#9806S, CST) with 1 mM PMSF (P7626, MilliporeSigma), and Phosphatase Inhibitor Cocktail 2 (P5726, MilliporeSigma). After quantifying total protein concentration with the BCA assay (23225, ThermoFisher), 10 µg of each sample was boiled at 95°C for 5 min and loaded onto the 4-12% Bis-Tris Gel (NP0336, ThermoFisher). Gels were run for 1 hour and 15 min at 120 V in with NuPage MES Buffer (NP0002, ThermoFisher). Proteins were then transferred onto polyvinylidene difluoride membrane (IPVH15150, MilliporeSigma) for 1 hour at 30 V in NuPage Transfer Buffer (NP0006, ThermoFisher). Membranes were checked for successful and adequate transfer with a Ponceau S stain (P7170, MilliporeSigma). Primary antibodies were diluted 1:1,000 (1 ug/mL) in blocking buffer (927-40000, Licor), and secondary antibodies were diluted to 1:10,000 (100 ng/mL) in 2.5% BSA (0332, VWR) in Tris buffered saline tween. All antibody incubations were performed overnight at 4°C. The primary antibodies used were rabbit antibody #9102L (CST) for ERK1/2, rabbit antibody #4370 (CST) for pERK, mouse antibody #31F12 (CST) for ALK in its kinase domain, rabbit antibody #Y1640 (CST) for pALK, and rabbit antibody #13E5 (CST) for *β*Actin. Anti-rabbit IgG #7074S (CST) and anti-mouse IgG #7076S (CST) secondary antibodies were used to detect the primary antibodies. Chemiluminescent signal was generated by SuperSignal West Pico PLUS Chemiluminescence substrate (345777, ThermoFisher), and detected on a BioRad ChemiDoc Imaging System. Membranes were stripped (21059, ThermoFisher) for 30 min, and re-blocked for 1 hour at room temperature.

Note on reproducibility: All of the analyses in this paper were performed in R v3.5.2. GraphPad prism v8 was used for dose response curve analysis. Amazon Web Services were used for the simulations in Fig. 1 and Supplementary Fig. 2. Version controlled html Rmarkdown files that include descriptions of the methods for all figures may be accessed via <u>Github repository</u>. This includes the code for the analysis of mutual exclusivity of gene pairs using pairwise comparisons of conditional selection. The results of all statistical analyses mentioned in this figure are included in the code. The pairwise comparisons workflow, outlined in our Github repository, requires an input dataset in the form of a table with columns for data on a gene of interest, and two positive control genes. We have also made available a function, named tcga_skcm_data_parser.Rmd, that can generate estimated counts for a gene of interest based on filters applied to RSEM, RPKM, and count data. The entire workflow was tested for reproducibility on simulated datasets and on expression data in LUAD for ALK and EGFR from the TCGA (41). The pairwise comparisons website, with the reproducible, interactive analyses, was created with the help of the r package WorkflowR (42).

## 9. Acknowledgements

We would like to thank Scott Leighow for his contribution with idea generation, critical analysis, and proofreading of this manuscript. We would also like to thank Kelly Hartsough, Lauren Randolph, Kyle Mcllroy, and Joshua Reynolds for their help revising previous versions of this manuscript. This publication was supported, in part, by NIH Grant T32GM108563.

---

**Supplementary figure 1:**
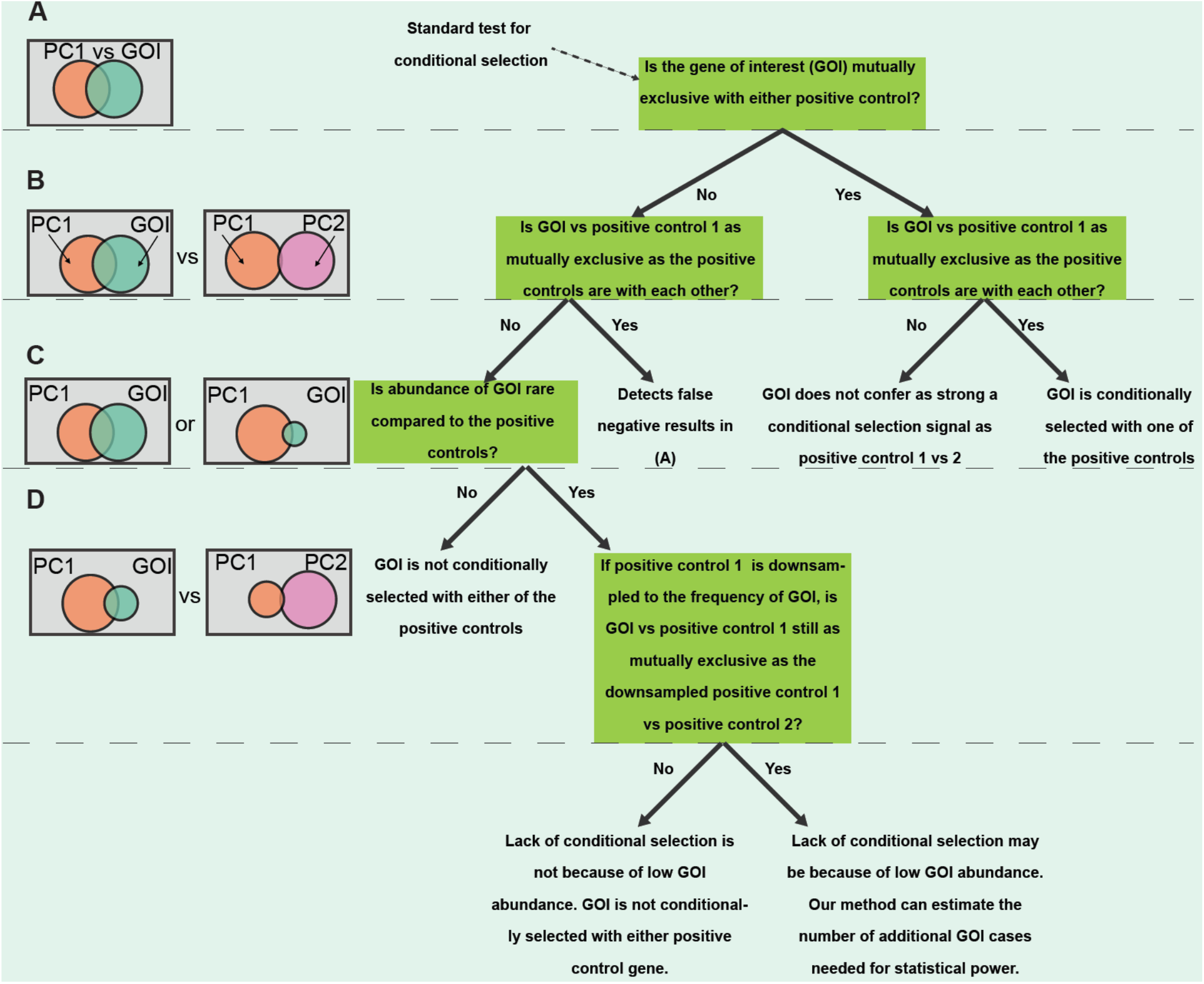
Decision tree for conditional selection analysis of rare variants. **(A)** A canonical test for mutual exclusivity compares a gene of interest to a positive control gene. **(B)** Pairwise comparisons enable comparing the gene pair in (A) to a previously established mutually exclusive gene pair. **(C)** If a gene of interest is rare compared to the positive controls in a cohort, statistical power is needed to conclude that the lack of signal is not because of low abundance. **(D)** Our method includes resampling simulations that downsample the positive control population to match the abundance of the gene of interest. If the abundance of the gene of interest is too rare for a fair comparison, our method determines the additional cases of GOI to perform a statistically significant comparison.

**Supplementary figure 2:**
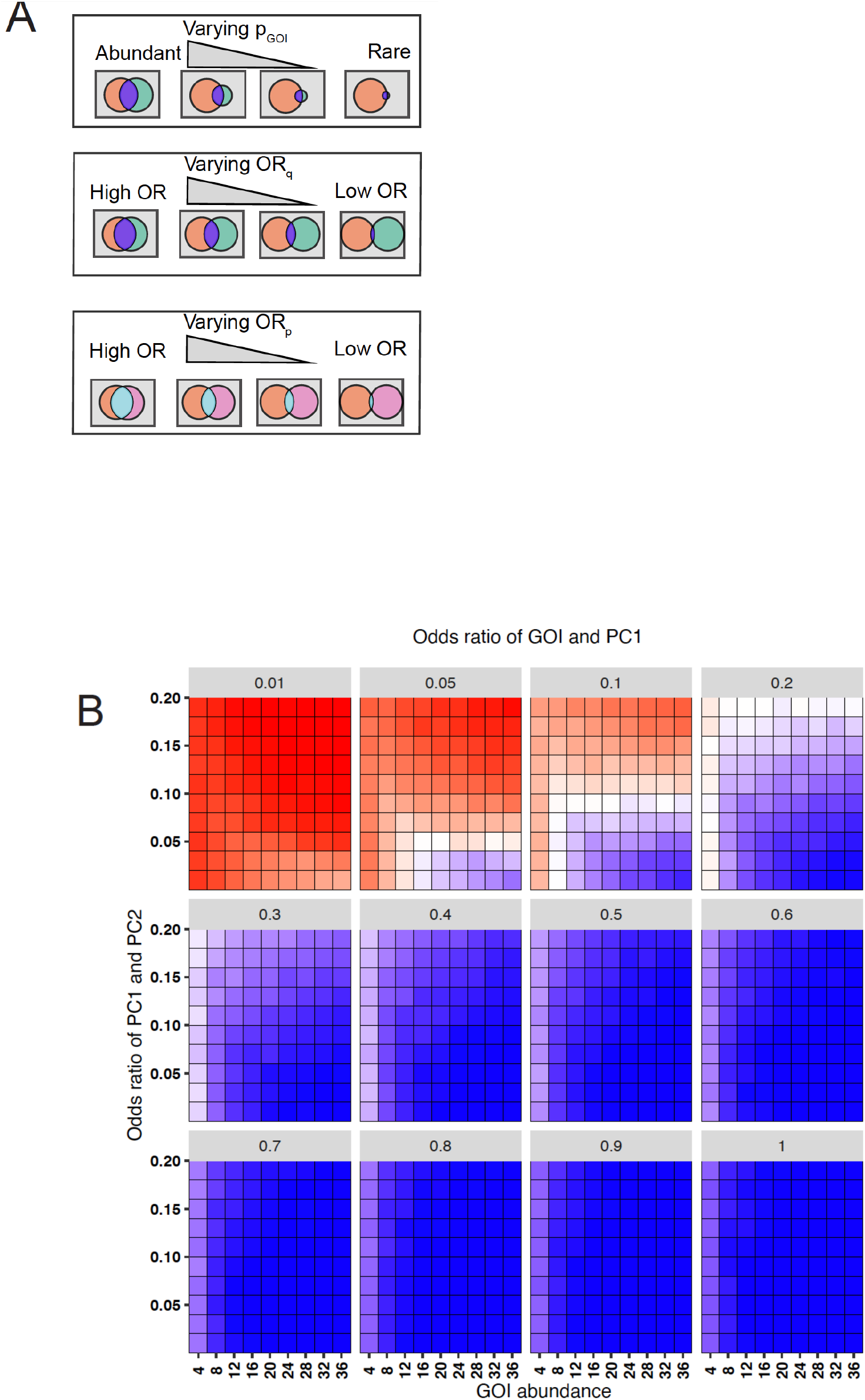
Pairwise comparisons of conditional selection in simulated cohorts. (**A)** Schematic of the different parameters that were varied when generating simulated cohorts with the gene of interest (teal), positive control 1 gene (orange), and the positive control 2 gene (pink). **(B)**. The minimum required abundance of the GOI to attain a high score decreases as the difference in the odds ratios between the two gene pairs increases. The positive control genes were simulated at moderate to high mutual exclusivity (OR of 0.01-0.2) while the GOI and PC1 were simulated at a range of mutual exclusivities (ORs between 0.01 and 1). A 500-patient cohort is used here but cohorts of 100-1,000 patients were tested, yielding similar results (not shown here).

**Supplementary figure 3:**
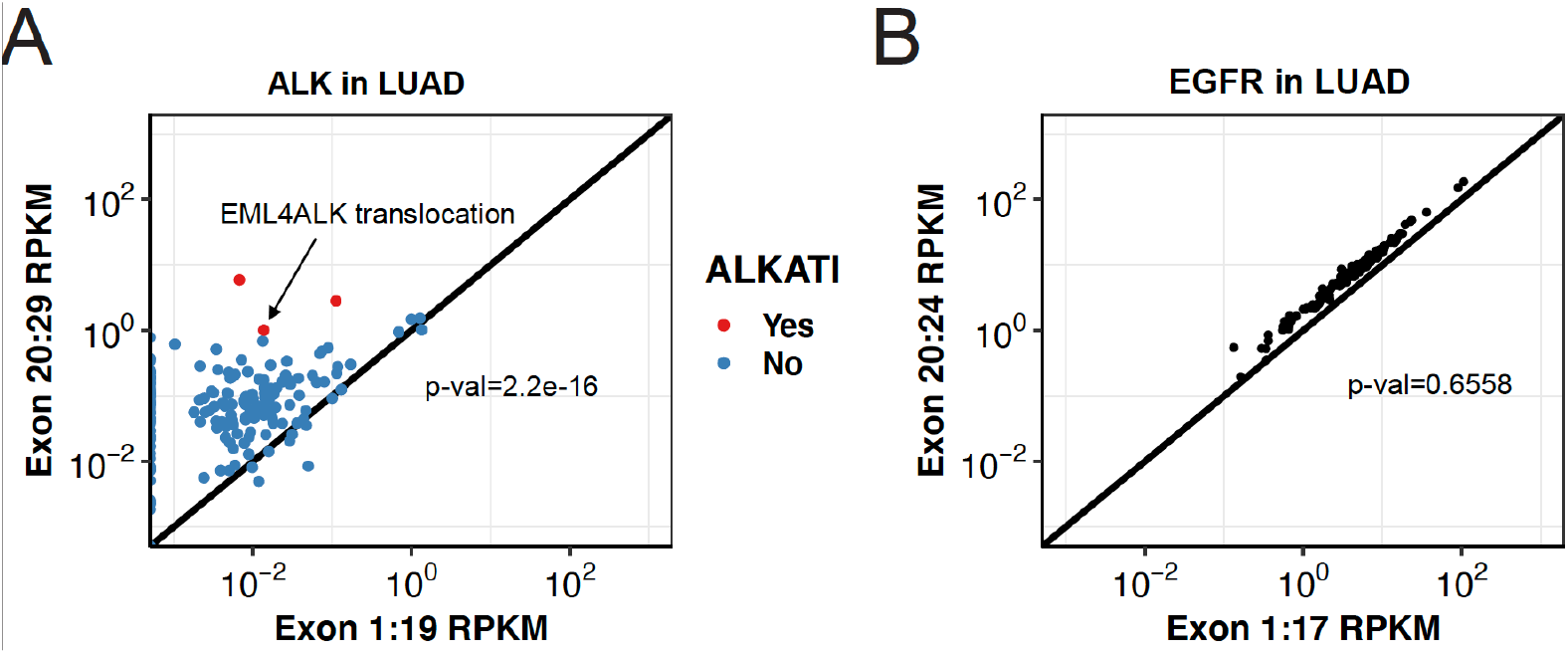
Propensity in expression towards kinase domain is ubiquitous in ALK. **(A)**. Expression profile of ALK^ATI^ patients in LUAD showing an exon imbalance towards the kinase domain (ex20-29). One of the patients matching ALK^ATI^ expression filters had an EML4ALK translocation. **(B)** Expression profile of EGFR in LUAD showing a lack of exon imbalance towards EGFR’s kinase domain (ex20-24).

**Supplementary figure 4:**
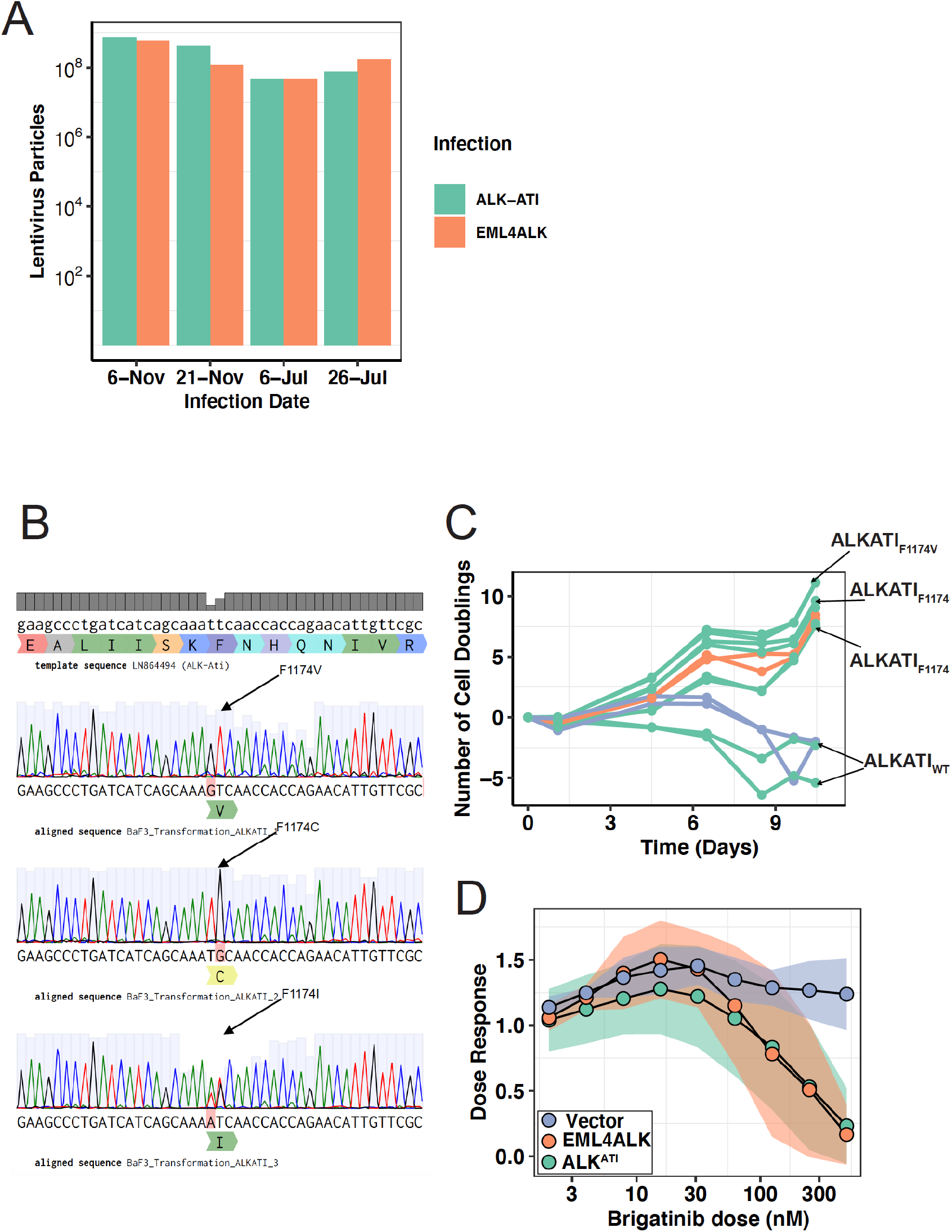
The confounding effect of transforming mutants on ALK^ATI^ transformation. **(A)** Counts of lentiviral particles across multiple infection attempts. Virus samples were collected prior to transducing Ba/F3s and SKML cell lines. **(B)** Nucleotide sanger sequencing of a part of the kinase domain of ALK showing the F1174C, V, and I transforming mutations. The three missense mutations represent the only mismatch in a conserved kinase domain. **(C)** Time to growth factor independence, as measured by number of cell doublings, for EML4-ALK, ALKATI^WT^, ALKATI^F1174C^, ALKATI^F1174V^, ALKATI^F1174I^, and vector control. **(D)** Brigatinib dose-response study for IL3 independent EML4-ALK, ALK^ATI^, and IL3 dependent vector-transduced Ba/F3 cells. Data are mean ± 95% confidence intervals (shaded region). 3, 5, and 1 independent transductions for EML4-ALK, ALK^ATI^, and vector control were tested. For all cell lines, 3 technical replicates were used at each concentration, and each assay was repeated on 3 separate days.

**Supplementary figure 5:**
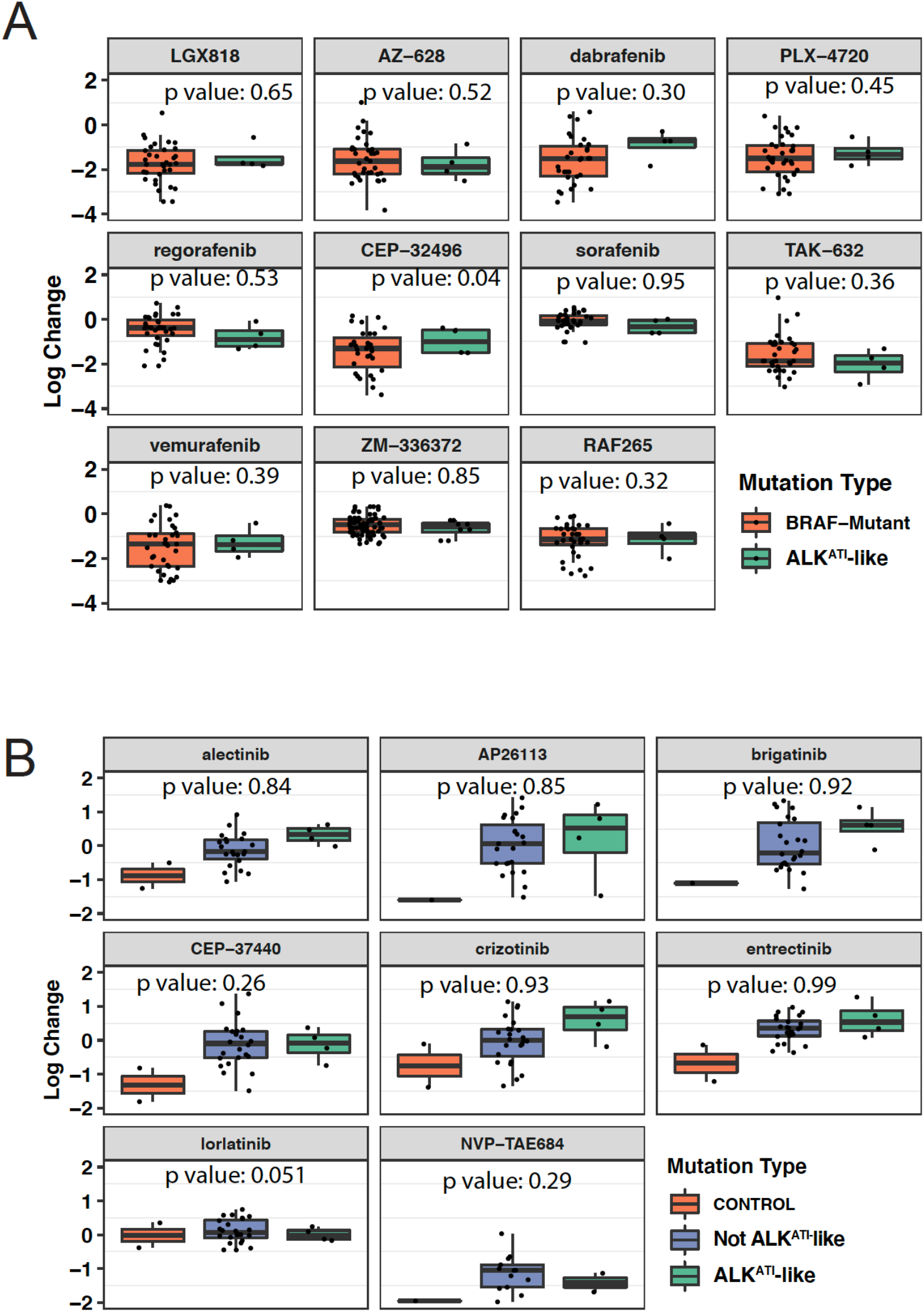
The presence of ALK^ATI^ does not predict an improved transforming potential or ALK-inhibitor sensitivity in melanoma. **(A)** Comparing the sensitivity to BRAF-inhibitors (n=11) of BRAF-mutated cell lines (orange, n=43), and BRAF-mutated cell lines that have an ALK^ATI^-like expression (cyan, n=4). P-value is from a one-sided multiple linear regression of inhibitor intolerance mapped as a function of ALK RSEM, RPKM, and Exon expression. **(B)** ALK-inhibitor sensitivity of melanoma cell lines with ALK^ATI^-like expression (cyan, n=4) or not (purple, n=32). Crizotinib sensitive control cell lines (orange) are Kelly (ALK^F1174L^ neuroblastoma), and NCI-H228 (EML4-ALK in lung adenocarcinoma). P-value is from a one-sided multiple linear regression of crizotinib sensitivity mapped as a function of ALK RSEM, RPKM, and Exon expression.

**Supplementary Table 1.** Multiple attempts to transduce oncogenes into BRAF_V600E_ melanoma lines fail. + represents outgrowth and – represents failure to grow for a transduction attempt. *Note: Replicates represent different infection attempts (biological replicates). All technical replicates for a given infection behaved similarly.

| Cell Line | SKMEL28 | G361 | SKMEL28 | G361 | SKMEL28 | G361 | BAF3 | HEK293 |
| --- | --- | --- | --- | --- | --- | --- | --- | --- |
| Plasmid | Vector | Vector | ALK<sup>ATI</sup> | ALK<sup>ATI</sup> | EML4ALK | EML4ALK | EML4ALK | EML4ALK |
| Selection | Puro | Puro | Puro | Puro | Puro | Puro | - iL3 | Puro |
| Replicate 1* | + | + | + | + | - | - | + | + |
| Replicate 2* | + | + | + | + | - | - | + | + |
| Replicate 3* | + | + | + | + | - | - | + | + |

**Supplementary Table 2.**
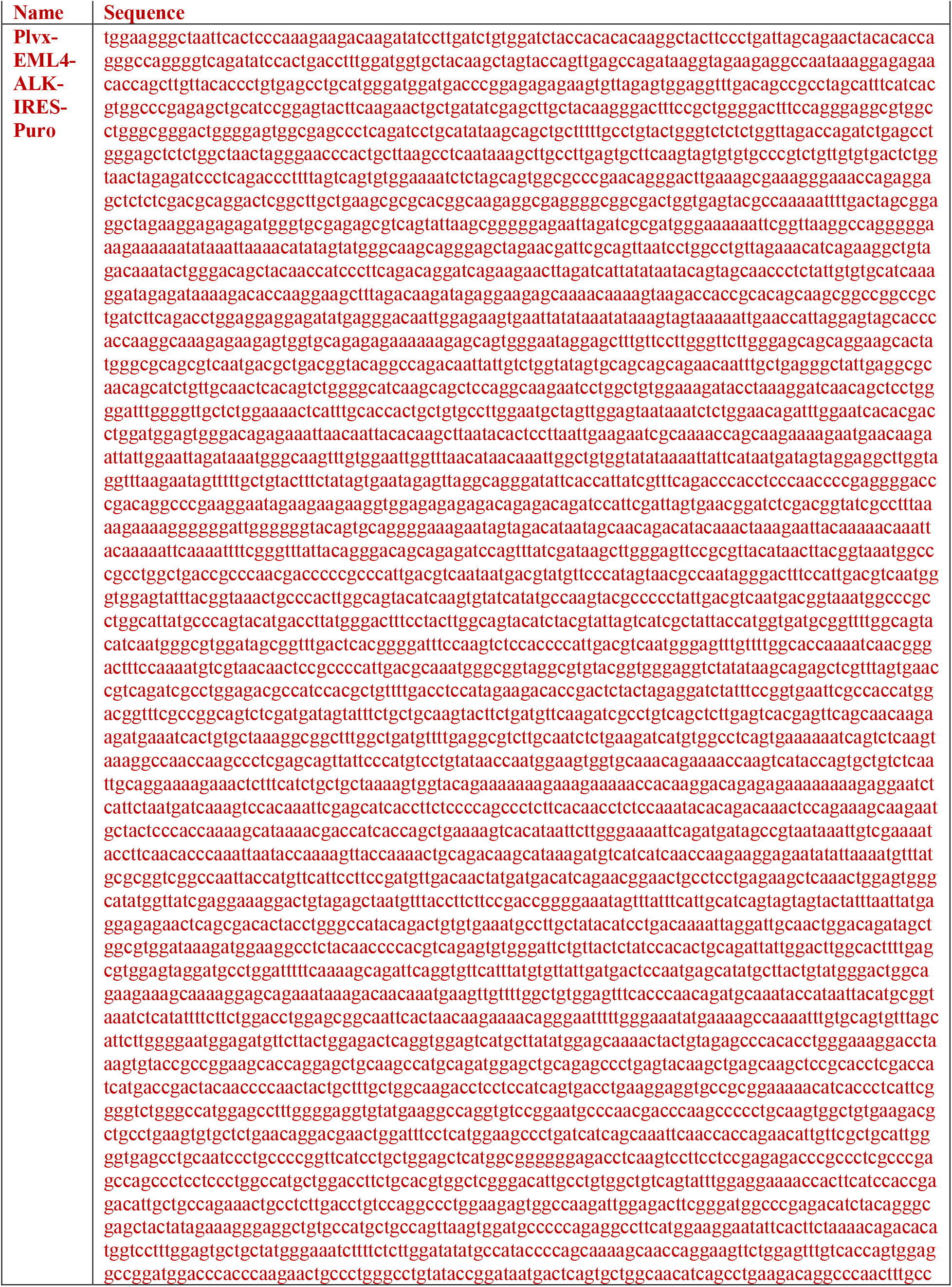
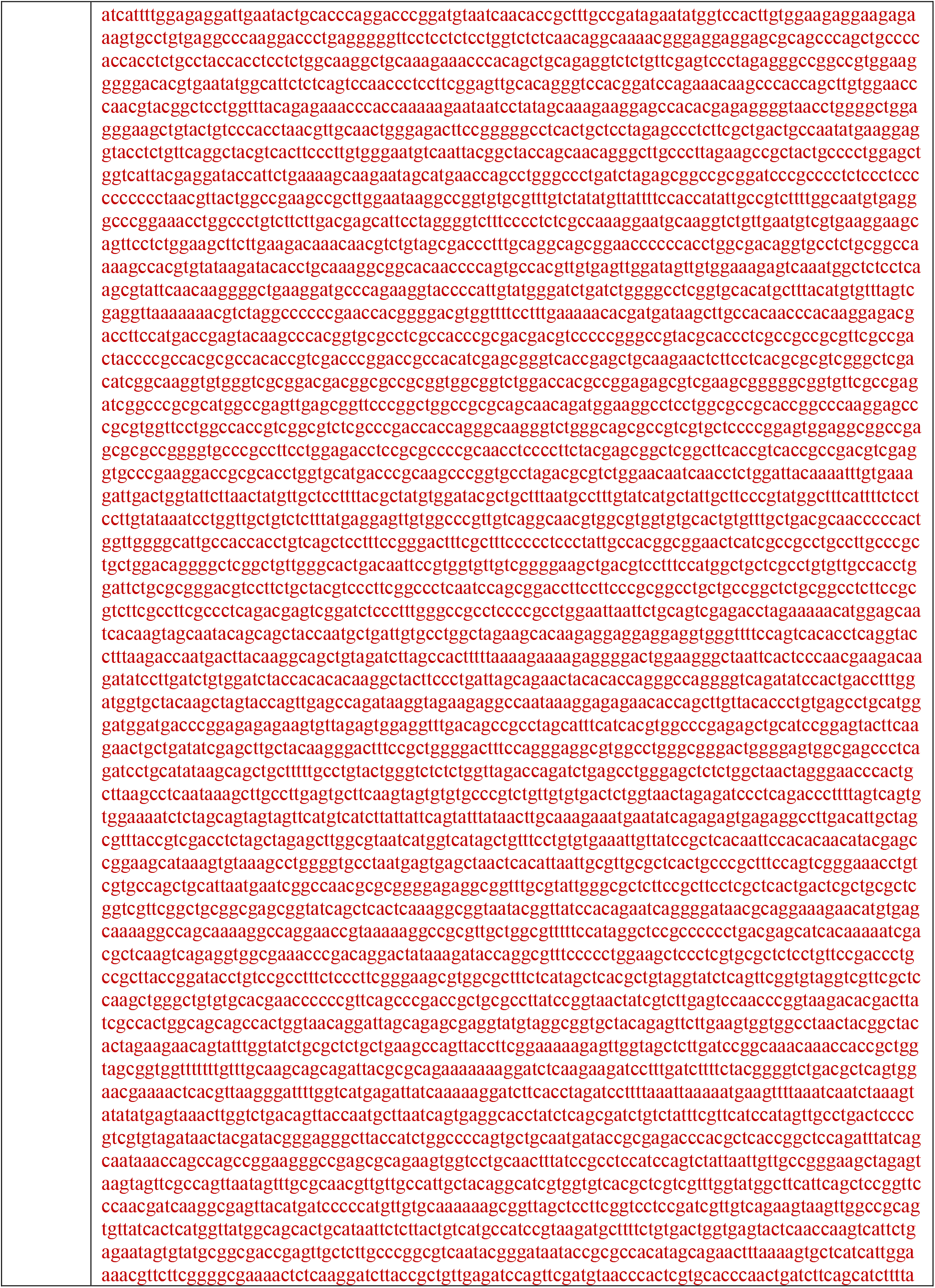
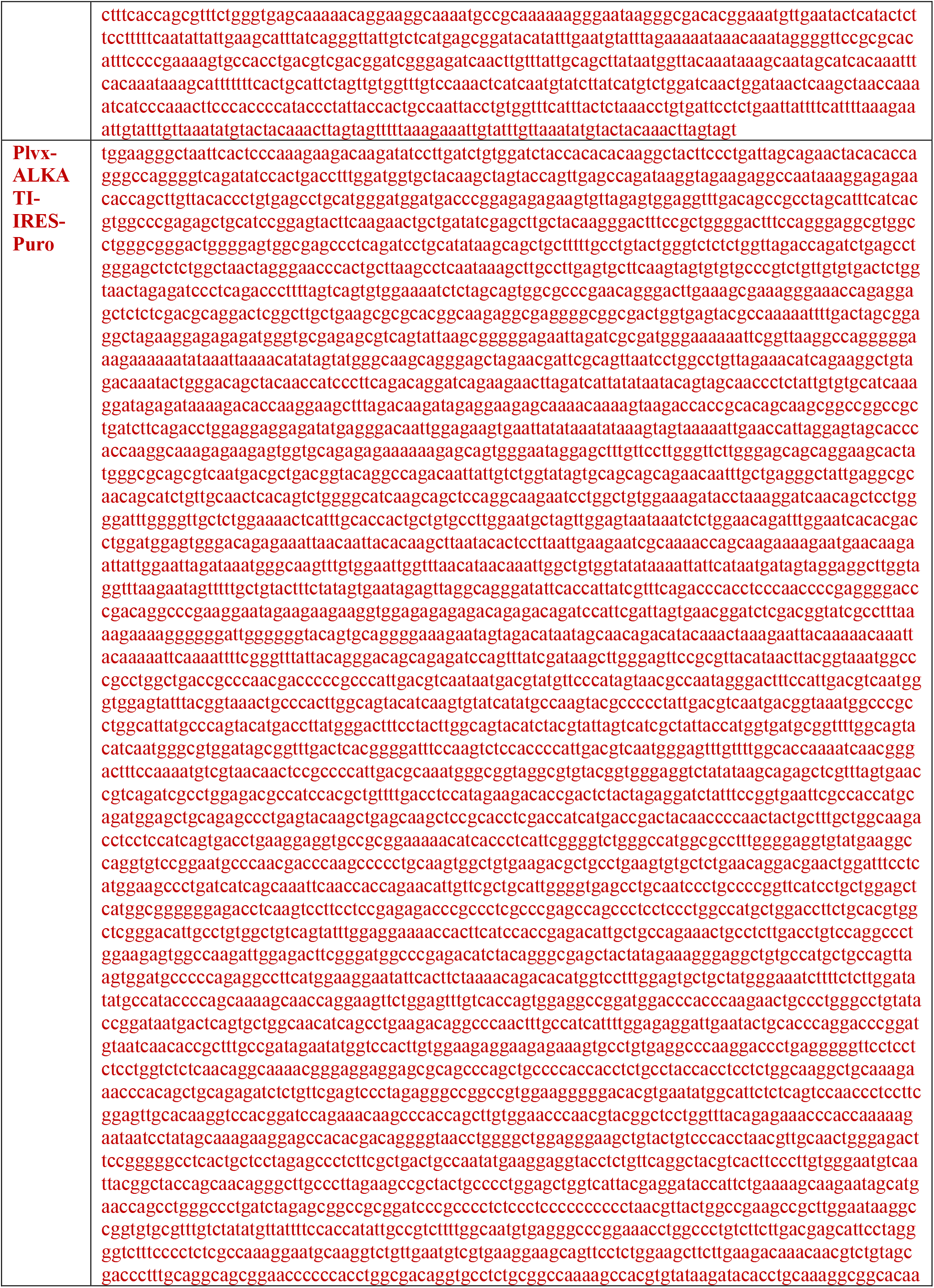
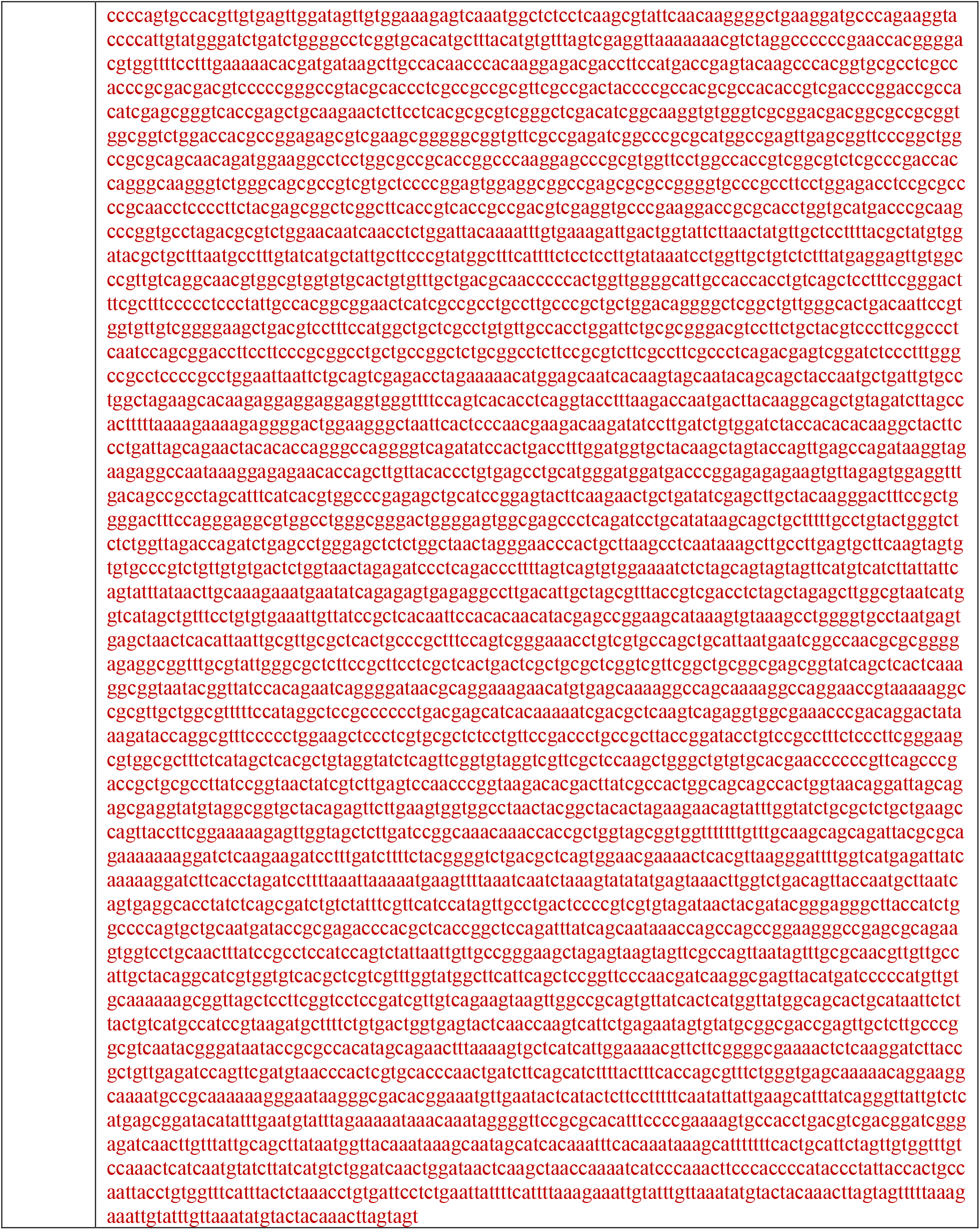
Sequence of lentiviral plasmid vectors with the associated transgene.

**Supporting Appendix S1: Pairwise comparisons pseudo-code**. The attached file contains pseudo-code with algorithms that detail the workflow for pairwise comparisons.

